# Compromised DNA replication in gut cells underlies tardigrade sensitivity to genotoxic stress

**DOI:** 10.1101/2025.07.23.666276

**Authors:** Gonzalo Quiroga-Artigas, Pauline Fontanié, Benjamin Lacroix, Maria Dolores Molina, María Moriel-Carretero

## Abstract

Tardigrades withstand severe DNA insults, including extreme doses of ionizing radiation, through unique protective proteins and strong upregulation of canonical DNA repair pathways. Yet, these extremophile animals are not immortal, and the cellular and organismal processes that ultimately fail under sustained genotoxic stress have not been characterized. Here, we identify DNA replication as the key vulnerability in the tardigrade *Hypsibius exemplaris*. Using the radiomimetic drug zeocin to induce DNA breaks, we show that continuous exposure progressively kills tardigrades, accompanied by striking body shrinkage and lipid depletion. DNA synthesis labeling reveals that zeocin disrupts replication and triggers *de novo* reparative synthesis in select non-dividing tissues. Pulse–wash experiments demonstrate that even transient damage to dividing gut cells irreversibly exhausts their replicative capacity, leading to midgut failure and animal death, despite systemic induction of DNA repair genes. Germ cells and embryos, with their high proliferation rates, show heightened sensitivity. Cross-phyla survival assays in the eutelic nematode *Caenorhabditis elegans* and neoblast-rich planarian *Schmidtea mediterranea* further link proliferative activity to mortality kinetics under DNA damage. Collectively, our findings pinpoint DNA replication as an Achilles’ heel of organismal survival under genotoxic stress, even in animals renowned for their extraordinary DNA damage tolerance.

## Introduction

DNA integrity is essential for organismal health and longevity: accumulated DNA damage drives mutagenesis, cancer, developmental defects, and aging (*1*). While most metazoans succumb to ionizing radiation (IR) at relatively low doses, tardigrades— microscopic, water-dwelling ecdysozoans—survive extreme genotoxic insults, including IR doses up to ∼5 kGy (*2*, *3*). This extremotolerance has sparked intense interest in tardigrades as models for space biology (*4*) and as sources of novel DNA-protection strategies with potential applications in medicine and biotechnology (*5*).

Mechanistic studies over the past decade have begun to reveal how tardigrades withstand DNA damage. In 2016, Hashimoto et al. discovered Dsup, a tardigrade-specific chromatin-binding protein that reduces oxidative DNA breaks (*6*) by shielding nucleosomes (*7*). Subsequent work showed that, upon exposure to IR or the radiomimetic agent bleomycin, tardigrades dramatically upregulate canonical repair pathways—including homology-directed repair (HDR), non-homologous end joining (NHEJ), base excision repair (BER), and microhomology-mediated end joining (MMEJ)—and deploy additional tardigrade-unique factors (TDR1, TRID1) to enhance DNA repair (*8–11*). Horizontal gene transfers enhancing antioxidant defenses and PARP1-mediated repair also contribute to their resilience (*10*). Yet, despite these formidable defenses, tardigrades are not immortal: the limits of their survival under sustained genotoxic stress, and the cellular and organismal processes that ultimately fail, have not been disclosed.

Until recently, tardigrades were thought to lack post-embryonic somatic cell proliferation, achieving eutely (fixed cell number) in adulthood. However, we and others have shown that *Hypsibius exemplaris*—our tardigrade model species (Fig. 1A)—retains two somatic populations that still divide after hatching (Fig. 1A; Fig. S1): storage cells, lipid-rich cells floating freely in the hemolymph, which replicate with each cuticle molt (*12*), and gut cells, which undergo continuous renewal from the anterior and posterior midgut (*13*). It has been proposed that organisms can endure high levels of DNA damage when their affected cells are non-proliferative or when proliferation plays a minimal role in maintaining organismal survival (*14*). This could be because the obligate progression through the cell cycle toward DNA replication represents a window of particular genomic vulnerability, as DNA damage can stall replication forks and induce further breaks (*15*), compromising cell viability and tissue integrity. In light of the revised understanding that tardigrades are not fully eutelic, the potential impact of compromised DNA replication on their survival under genotoxic stress warrants investigation.

**Figure 1.**
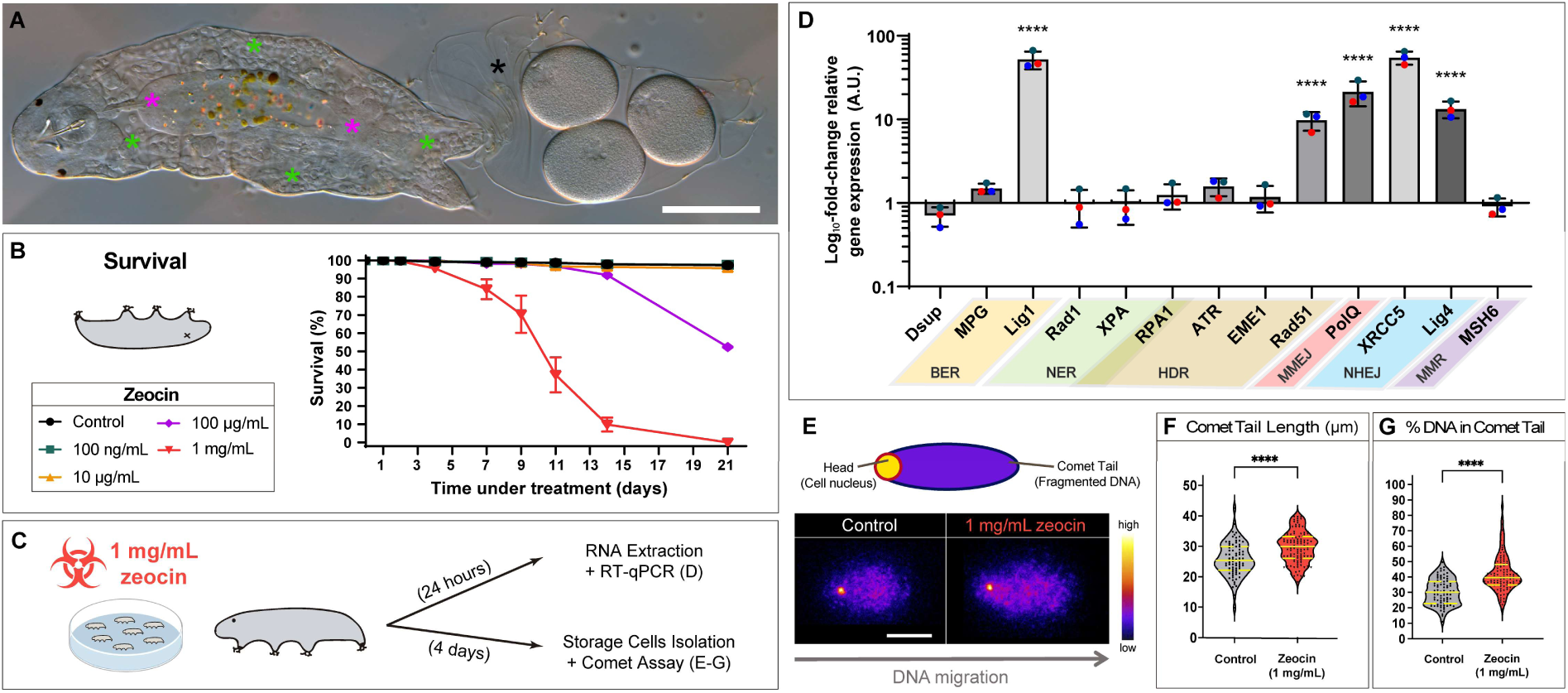
Zeocin causes DNA damage and reduces survival in the tardigrade *Hypsibius exemplaris*. (A) Representative image of *H. exemplaris* following a molting and egg-laying event. Green asterisks indicate storage cells throughout the body cavity; magenta asterisks mark the foregut and hindgut; black asterisk marks the molted exuvia containing three eggs. (B) Survival of adult *H. exemplaris* under increasing zeocin concentrations. Data are shown as mean ± standard error of the mean (SEM) from three to seven biological replicates, each consisting of 60–120 animals. (C) Schematic illustrating the experimental design related to panels D–G. (D) RT-qPCR analysis showing log_10_-transformed relative gene expression levels of *Dsup* and several DNA repair pathway genes in animals treated with 1 mg/mL zeocin for 24 hours, compared to untreated controls. Bar heights represent the mean of three independent experiments; error bars indicate standard deviation. Each colored circle corresponds to one independent experiment. A.U. = arbitrary units. Pathway abbreviations: BER, base excision repair; NER, nucleotide excision repair; HDR, homology-directed repair; MMEJ, microhomology-mediated end joining; NHEJ, non-homologous end joining; MMR, mismatch repair. (E) Schematic of comet assay readout (top). Representative fluorescence microscopy images of storage cell nuclei from control and zeocin-treated animals (bottom). DNA migration direction is indicated by an arrow. Images are pseudo-colored using the ‘Fire’ LUT in ImageJ; the adjacent color scale reflects relative pixel intensity. (F) Violin plots showing comet tail length measurements. (G) Violin plots showing the percentage of DNA in the comet tail. For panels F–G, experimental conditions match those in panel E. Yellow solid lines represent median values; dashed yellow lines indicate quartiles. Violin width reflects data point density. Individual data points are shown as circles (control) and triangles (zeocin-treated). *n* = 79 (control), *n* = 104 (zeocin-treated). **** *p* ≤ 0.0001. Scale bars = 50 μm in A; 20 μm in E.

In this study, we exposed adult *H. exemplaris* to the radiomimetic agent zeocin—a bleomycin-family drug inducing single- and double-strand DNA breaks (*16*)—and examined the organism-wide effects of DNA damage. We confirmed that zeocin remains active in the tardigrade growth medium, elicits robust induction of DNA repair genes, and causes extensive DNA fragmentation. Notably, zeocin induced a dramatic reduction in body size and depletion of fat stores. We found that it disrupts gut cell replication, likely leading to midgut failure, nutritional collapse, and ultimately death. In addition, we detected *de novo* reparative DNA synthesis in select non-dividing somatic tissues, consistent with HDR occurring only in a subset of tardigrade cell types. Pulse–wash experiments of increasing duration further implicated the irreversible exhaustion of gut cell replicative capacity as the primary driver of tardigrade mortality. Thus, DNA replication emerges as the central vulnerability that ultimately limits survival under genotoxic stress in these extremotolerant animals. Finally, we reinforced this notion by showing that germ cells and embryos are particularly sensitive to zeocin, consistent with their elevated replication rates.

In sum, despite their remarkable arsenal for repairing and tolerating DNA damage, tardigrades are fundamentally constrained by the need to safely replicate their genomes. This work establishes a foundation for broader exploration of how replication-associated vulnerabilities shape organismal responses to genotoxic stress.

## Results

### The radiomimetic agent zeocin induces DNA damage and causes tardigrade death

We first evaluated tardigrade survival under continuous zeocin exposure at different concentrations (Fig. 1B). To verify that zeocin retained its activity over time, we incubated *S. cerevisiae* yeast cells for 3 hours in the medium where tardigrades had been exposed to zeocin for 7 days. Using western blotting, we assessed activation of the DNA damage response *via* Rad53 phosphorylation (*17*). This assay confirmed that zeocin remained active after 7 days in the tardigrade medium (Fig. S2), and we thus refreshed the medium weekly in all long-term experiments. While 100 ng/mL and 10 μg/mL had no measurable effect on survival, 100 μg/mL increased long-term mortality. At 1 mg/mL, zeocin began to reduce survival by day 4, with a steep decline between days 4 and 14 (from ∼95% to ∼10%), followed by near-complete mortality by day 21 (Fig. 1B). These results show that, although *H. exemplaris* is notably resistant to zeocin, it cannot withstand sustained exposure to high levels of genotoxic stress indefinitely.

Next, to confirm that zeocin effectively reaches internal tissues and induces DNA damage, we exposed adult animals to 1 mg/mL zeocin for 24 hours and extracted total RNA for RT-qPCR analysis (Fig. 1C). We profiled a panel of evolutionarily conserved DNA repair genes spanning all major pathways (Fig. 1D). In line with recent studies using IR and bleomycin (*8*, *9*), *H. exemplaris* significantly upregulated several DNA repair genes, mostly related to double-strand break (DSB) repair—*Lig1*, *Rad51*, *PolQ*, *XRCC5*, and *Lig4* (Fig. 1D). This response suggests that zeocin induces substantial DNA damage, likely both single- and double- strand breaks. We also examined the expression of the tardigrade-specific gene *Dsup*, known to confer DNA damage protection (*6*), and found no significant changes (Fig. 1D), consistent with prior RNA-seq datasets from irradiated animals (*8*, *9*). To further validate that zeocin causes DNA damage in *H. exemplaris*, we treated adults with 1 mg/mL zeocin for four days—coinciding with the initial decline in survival (Fig. 1B)—and performed comet assays on isolated storage cells (Fig. 1C). The comet assay, which detects DNA strand breaks at the single-cell level (*18*, *19*), and has been previously applied to tardigrade storage cells (*20*), revealed clear evidence of fragmentation: zeocin-treated nuclei exhibited significantly longer comet tails and higher DNA content in the tails compared to controls (Fig. 1E–G; Fig. S3), confirming zeocin’s genotoxicity. These results provided a framework to explore the cellular and physiological mechanisms underlying tardigrade mortality.

### Zeocin reduces body size, depletes fat reservoirs, and alters cuticle structure

In long-term incubations, zeocin-treated animals consistently appeared visibly smaller and exhibited empty or partially empty guts (Fig. 2A). To quantify this shrinkage, we measured the body length of grown adults (≥240 μm) after 10 days of exposure to 1 mg/mL zeocin, and compared it to the effects of starvation in parallel. While starvation alone caused a significant reduction in body size relative to controls, the size decrease was even more pronounced in zeocin-treated animals (Fig. 2A, B).

**Figure 2.**
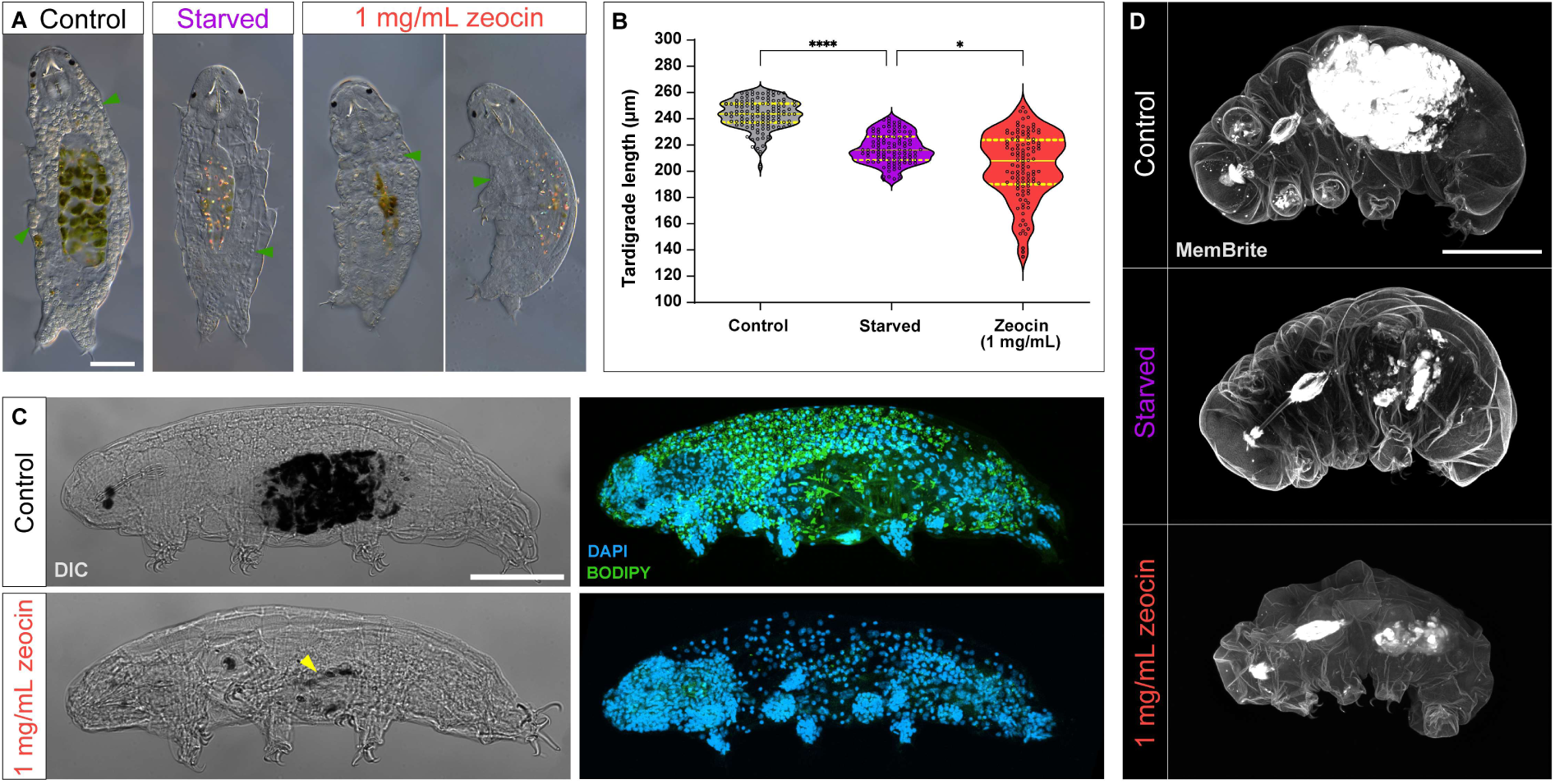
Zeocin exposure leads to reduced animal size, lipid loss, and cuticle alterations. (A) Representative images of *H. exemplaris* following 10-day starvation or 1 mg/mL zeocin exposure. Green arrowheads point to storage cells. Note the empty guts in both conditions. (B) Violin plots showing body length of tardigrades under different conditions — exposed for 10 days (*n* = 121 [control], *n* = 105 [starved], *n* = 109 [zeocin-treated]). Individual measurements are shown as empty circles. Plot format as in Fig. 1F–G. (C) Representative images of *H. exemplaris* after 4 days in 1 mg/mL zeocin. Left: animal morphology (DIC, gray); digested algae appear black in the gut. Yellow arrowhead points to an empty gut in the zeocin- treated condition. Right: nuclei (DAPI, blue) and neutral lipids (BODIPY, green), mainly localized in lipid-rich storage cells. (D) Representative images of *H. exemplaris* following 10- day starvation or 1 mg/mL zeocin exposure, highlighting cuticular structures (MemBrite, gray). Algae within the gut auto-fluoresce in the same channel. Confocal images are maximum projections of full-depth z-stacks. **** *p* ≤ 0.0001; * *p* ≤ 0.05. Scale bars = 50 μm.

To understand this phenotype, we first tested whether the quality of the algal food source (*Chlorococcum sp*.) had been compromised by zeocin, potentially deterring ingestion. Algae were pre-treated with 1 mg/mL zeocin for 48 h, then washed and fed to 24-h–starved, untreated animals. Within 24 h, all animals had consumed the zeocin-treated algae (Fig. S4), ruling out poor algal quality as a feeding deterrent. We then asked whether extended zeocin exposure impaired the animals’ ability to feed. Animals pre-treated with zeocin for 7 days failed to ingest fresh algae, whereas 7-day–starved controls did so normally (Fig. S5). Consistently, lipid reservoirs were dramatically depleted in zeocin-treated animals after 4 days, as revealed by BODIPY staining (Fig. 2C), paralleling the effect observed under starvation (*12*). These results suggest that zeocin damages the digestive system, preventing food intake, likely contributing to reduced body size and survival.

Notably, although zeocin-treated and starved animals exhibited similarly reduced sizes—with much of the size distribution overlapping (Fig. 2B, compare violet and red plots)— a subset of zeocin-treated individuals appeared even smaller. We hypothesized that additional tissue-level damage might contribute to this residual phenotype. The cuticle, synthesized by epidermal cells, is a key structure in tardigrades that supports environmental resilience and body integrity (*21*). To assess whether cuticle structure was affected, we stained live animals with MemBrite, which labels the cuticle, buccopharyngeal apparatus, and claws. Zeocin-treated animals appeared not only smaller than controls and starved animals, but also showed markedly dimmer MemBrite fluorescence and disrupted cuticle folding (Fig. 2D). To investigate potential cellular causes, we examined the nuclei of epidermal cells after one week of treatment using DAPI staining and confocal microscopy. Epidermal nuclei in treated animals displayed signs of fragmentation (Fig. S6), indicating that DNA damage compromises epidermal integrity. Together, these observations suggest that zeocin impairs both the structure and function of the epidermis and cuticle, along with the digestive system, synergistically driving physiological decline.

### Zeocin impairs DNA replication and triggers repair responses across tissues

To uncover molecular indicators that might explain the observations presented above, we incubated animals with 5-ethynyl-2′-deoxyuridine (EdU), a thymidine analog that labels newly synthesized DNA. This approach is based on the principle that EdU gets incorporated into DNA during both replication and repair. Among repair mechanisms, HDR is especially likely to incorporate substantial EdU, as it involves synthesis of long DNA stretches using a homologous template (*22*). EdU incorporation has been successfully used to monitor DNA replication in tardigrades (*12*, *13*), and EdU-based detection of DNA repair has been demonstrated in the rotifer *Adineta vaga* (*23*). Thus, EdU labeling enables the mapping of these two DNA-related processes—or their absence—across different tardigrade tissues.

In fully-grown adult tardigrades, the main somatic cell type that replicates are the gut cells, which do so continuously (*13*). In contrast, the other major replicative somatic cell type, the storage cell, exhibits only residual levels of replication, as most replication events are coupled to molting episodes during growth (*12*). We co-incubated adult animals with 1 mg/mL zeocin and EdU for 4 days (Fig. 3A) and made several key observations. First, zeocin significantly reduced the number of EdU⁺ gut cells compared to untreated controls (Fig. 3B, E), revealing that DNA damage impairs gut cells DNA replication in *H. exemplaris*. To test whether replication in storage cells is similarly affected, we turned to molting-stage juveniles, where storage cells actively replicate. Juveniles were incubated with 1 mg/mL zeocin and EdU for 2 days (Fig. 3C). We observed a significant reduction in EdU⁺ storage cells in treated juveniles compared to controls (Fig. 3D, E), confirming that zeocin also impairs replication in this cell type. To determine whether the lack of replication in storage cells was critical for survival, we hypothesized that juveniles would be more sensitive to zeocin than adults. However, survival assays under continuous zeocin exposure showed that juveniles exhibit similar resistance to adults (Fig. S7; Fig. 1B), suggesting that reduced storage cell replication does not drive mortality, and instead pointing to impaired replication in gut cells as a more likely contributor.

**Figure 3.**
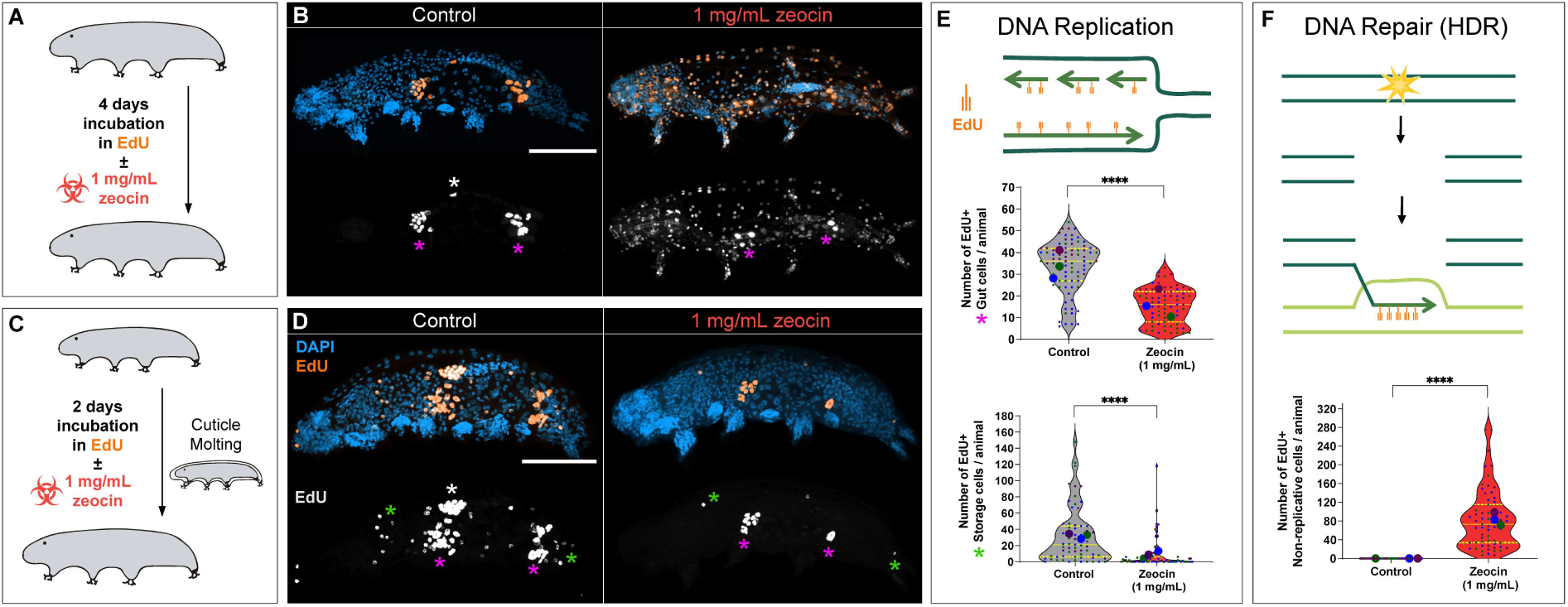
Zeocin impairs DNA replication in proliferative cells and activates DNA repair in various cell types. (A) Schematic of a 4-day EdU ± zeocin incubation experiment, corresponding to panels B, E (middle), and F. (B) Representative images of *H. exemplaris* after 4-day EdU ± zeocin exposure, showing nuclei (DAPI, blue) and EdU^+^ cells (orange). Bottom images display EdU^+^ signal only (gray). (C) Schematic depicting a 2-day EdU ± 1 mg/mL zeocin incubation experiment during a molting cycle, corresponding to panels D and E (bottom). (D) Representative images of *H. exemplaris* following 2-day EdU ± zeocin exposure during molting, showing nuclei (DAPI, blue) and EdU^+^ cells (orange). An example lacking EdU incorporation in non-replicative cells was selected to clearly illustrate the impairment of storage cell replication in zeocin-treated juveniles compared to controls. Bottom images display EdU^+^ signal only (gray). In panels B and D, green asterisks indicate EdU^+^ storage cells; magenta asterisks mark EdU^+^ gut cells in the foregut and hindgut; white asterisk marks EdU^+^ germ cells. (E) Top: schematic illustrating EdU incorporation during DNA replication. Middle: violin plots showing the number of EdU^+^ gut cells per animal (*n* = 79 [control], *n* = 73 [zeocin-treated]). Bottom: violin plots showing the number of EdU^+^ storage cells per animal (*n* = 62 [control], *n* = 69 [zeocin-treated]). Data are from three independent experiments. Each colored dot represents an individual animal; dot color indicates experiment; larger dots represent per-experiment means. Plot style as in Fig. 1F–G. (F) Top: schematic showing EdU incorporation during homologous-directed repair (HDR). Bottom: violin plots showing the number of EdU^+^ non-replicative cells per animal (*n* = 84 [control], *n* = 60 [zeocin-treated]). Plot details as in panel E. Images are maximum projections of full-depth z-stacks. **** *p* ≤ 0.0001. Scale bars = 50 μm.

The second key observation was that continuous zeocin exposure triggered EdU incorporation in cell types not typically associated with replication—such as claw gland cells, epidermal cells, and cells in the brain and pharynx (Fig. 3B, F). However, not all non-replicative cell types incorporated EdU under these conditions; notably, ganglion cells and trophocytes did not (Fig. 3B; Fig. S8). Storage cells did not incorporate EdU either (Fig. 3B; Fig. S8). To distinguish between replication- and repair-driven EdU incorporation, we quantified fluorescence intensity in EdU⁺ non-replicative cells and compared it to that in replicative gut and storage cells. Because repair synthesis involves only DNA tracts, EdU signal is expected to be weaker than during genome-wide replication. Indeed, EdU fluorescence was significantly lower in non-replicative cells (Fig. S9), consistent with DNA synthesis during repair. Notably, this pattern aligns with the upregulation of *RAD51*, a key HDR factor, in zeocin-treated animals (Fig. 1D). Altogether, these findings suggest that zeocin-induced DNA damage impairs replication in proliferative cell types, while simultaneously triggering *de novo* DNA synthesis associated with repair—likely through the HDR pathway— in various non-proliferative somatic tissues.

### The replicative ability of gut cells is exhausted upon DNA damage

The results presented so far suggest that loss of homeostasis in gut cells may be the primary driver of nutritional decline, size reduction, and diminished viability in zeocin-treated animals. Notably, the anterior foregut and posterior hindgut cells have been proposed to serve as midgut epithelial precursors, proliferating steadily to replenish aging gut cells (*13*, *24*).

To assess the long-term vulnerability of these gut cell populations, we exposed adult animals to 1 mg/mL zeocin for increasing durations (24 hours, 4 days, or 9 days), followed by drug removal and a recovery period. On day 9, all groups received a 24-hour EdU pulse to evaluate the replicative activity in the gut (Fig. 4A). We observed a clear, progressive decline in EdU⁺ gut cells with longer zeocin exposures (Fig. 4B, C). These results suggest that DNA damage irreversibly impairs gut cell replication, despite the otherwise robust repair capacity of tardigrades, and that the severity of this impairment increases with the duration of the DNA-damaging treatment.

**Figure 4.**
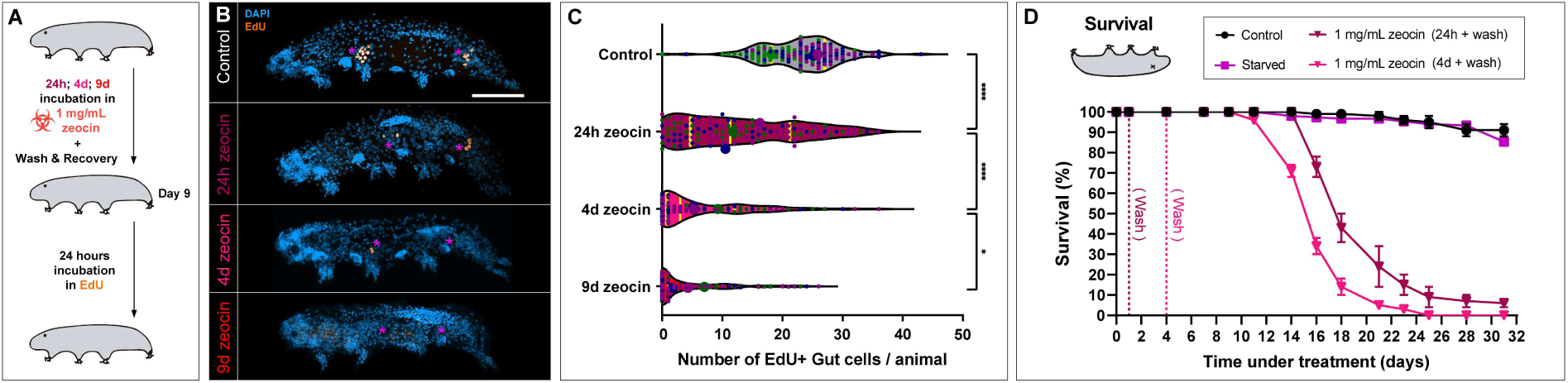
Zeocin exhausts the replicative capacity of gut cells, accelerating mortality. (A) Schematic illustrating the experimental design for panels B–C. Animals were exposed to 1 mg/mL zeocin for varying pulse durations (24 hours, 4 days, or 9 days), followed by a washout and recovery period. By day 9, all samples were incubated for 24 hours with EdU. (B) Representative images of *H. exemplaris* following different zeocin pulse durations, showing nuclei (DAPI, blue) and EdU⁺ cells (orange). Magenta asterisks mark the foregut and hindgut. All images are maximum intensity projections from full-depth z-stacks. Scale bar = 50 μm. (C) Violin plots showing the number of EdU⁺ gut cells per animal (*n* = 120 [control], *n* = 111 [zeocin 1 day], *n* = 107 [zeocin 4 days], *n* = 166 [zeocin 9 days]). Data represent three independent experiments. Each colored dot corresponds to an individual animal, with dot color indicating the experiment; larger dots show the per-experiment means. Plot style as in Fig. 1F–G. (D) Survival curves of adult *H. exemplaris* after starvation or zeocin pulse treatments (1 mg/mL), followed by washout. Data are shown as mean ± SEM from two to three biological replicates, each with 50 animals. **** *p* ≤ 0.0001; * *p* ≤ 0.05.

To understand whether this gut cell exhaustion could underlie the long-term physiological decline and mortality, adults were exposed to continuous starvation, or to 1 mg/mL zeocin for either 24 hours or 4 days, then washed and monitored for survival over one month (Fig. 4D). By day 31, more than 90% of control animals, and about 85% of starved animals remained alive. The big majority of surviving starved animals were able to feed again after one month of food deprivation. In contrast, a 24-hour zeocin pulse began reducing survival by day 16, with a sharp decline between days 16 and 25 (from ∼70% to ∼10%). Animals surviving beyond day 25 appeared healthy, resumed feeding, and laid eggs, suggesting that a small fraction could fully recover from the treatment. The 4-day zeocin pulse produced more severe effects: survival began to drop by day 11, fell precipitously between days 11 and 21 (from ∼95% to ∼5%), and reached 100% mortality by day 25 (Fig. 4D).

These findings demonstrate that even a transient exposure to zeocin causes irreversible DNA damage, leading to delayed mortality. Although animals survive longer after pulse treatment (Fig. 4D) than under continuous exposure (Fig. 1B), the damage ultimately results in irreversible gut failure. Altogether, our data show that zeocin exhausts the replicative potential of gut cells, in contrast to starvation (*12*, *13*), impairing epithelial renewal in the midgut, disrupting nutrient assimilation, and ultimately condemning animals to die in the mid- term.

### Embryonic sensitivity to zeocin underscores replication as a central vulnerability in tardigrades

The strong effects of zeocin on gut cell renewal prompted us to ask whether other replicative tissues or developmental stages might also be particularly susceptible to DNA damage. To explore this, we turned our attention to germ cells—which replicate in coordination with molting (*12*)—and embryos, which are highly proliferative (*25*).

Tardigrade germ cells generate both trophocytes and oocytes through mitotic division (*26*, *27*). Because oocytes ultimately give rise to parthenogenetic embryos in *H. exemplaris*, their genomic integrity is even more essential than in other sexually reproducing animals. To assess how DNA damage impacts germ cell function, we quantified fertility by counting the number of eggs laid per animal under continuous exposure to increasing zeocin concentrations. While 100 ng/mL and 10 µg/mL had no measurable effect, 100 µg/mL significantly reduced egg production, and 1 mg/mL induced complete sterility (Fig. 5A; Fig. S10). Pulsed exposures to 1 mg/mL zeocin also revealed duration-dependent effects: a 24- hour pulse substantially reduced fertility, while a 4-day pulse rendered animals sterile (Fig. 5B; Fig. S10). These results indicate that tardigrade germ cells are vulnerable to DNA damage in a dose- and time-dependent manner, consistent with previous findings using IR (*2*), and at lower doses than those required to compromise adult viability (Fig. 1B).

**Figure 5.**
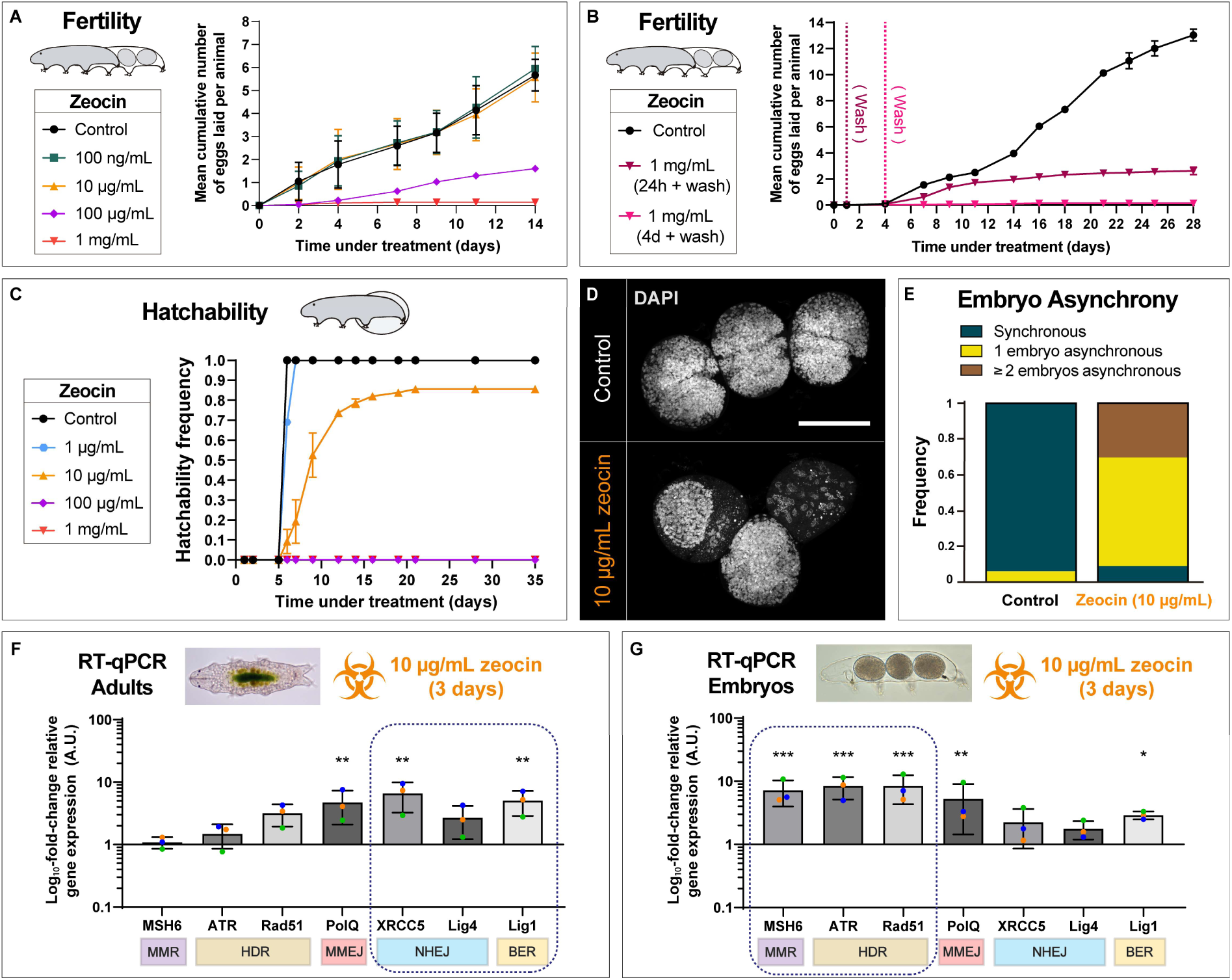
Zeocin impairs fertility, hatchability, and embryonic development in tardigrades. (A) Cumulative number of eggs laid per alive animal over 14 days of continuous zeocin treatment at varying doses. Data are shown as mean ± SEM from three biological replicates (60–120 animals each). (B) Cumulative number of eggs laid per alive animal over 28 days following pulsed zeocin treatments (24 h pulse + wash; 4-day pulse +W wash) at 1 mg/mL. Data are shown as mean ± SEM from two biological replicates (50 animals each). (C) Hatchability over time of *H. exemplaris* embryos continuously exposed to varying zeocin doses. Data are shown as mean ± SEM from three biological replicates (50–100 embryos each). (D) Representative confocal images of 3-day-old embryos stained with DAPI (white) under control and 10 μg/mL zeocin treatment. Each image shows three embryos per exuvia. Images are maximum z-projections. Scale bar = 50 μm. (E) Frequency of developmental asynchrony in embryos exposed to control or 10 μg/mL zeocin, scored at day 3. Two pooled biological replicates (*n* = 49 control; *n* = 63 zeocin). Only exuviae containing ≥3 embryos were analyzed. (F) RT-qPCR analysis showing log_10_-transformed relative gene expression levels of DNA repair genes in adults treated with 10 μg/mL zeocin for 3 days, compared to untreated controls. (G) RT-qPCR analysis showing log_10_-transformed relative gene expression levels of DNA repair genes in embryos treated with 10 μg/mL zeocin for 3 days, compared to untreated controls. In F–G, bars represent means of three independent experiments; error bars indicate standard deviation. Each colored circle corresponds to one experiment. Pathway abbreviations as in Fig. 1; outlined boxes highlight pathways differentially upregulated between adults and embryos. A.U. = arbitrary units. *** *p* ≤ 0.001; ** *p* ≤ 0.01; * *p* ≤ 0.05.

Next, we assessed how embryonic development responds to continuous zeocin exposure by monitoring hatchability (Fig. 5C). Gravid adults were incubated with the drug shortly before oviposition, ensuring that embryos were exposed from the earliest developmental stages. Untreated controls hatched after about six days, as previously reported (*2*). Embryos treated with 1 µg/mL hatched slightly later, while those treated with 100 µg/mL or 1 mg/mL failed to hatch altogether. Notably, at 10 µg/mL, ∼80% of embryos eventually hatched, albeit with pronounced delays—some requiring up to three weeks, compared to six days in untreated conditions (Fig. 5C). These results demonstrate that actively dividing embryos are more vulnerable to DNA damage than adult animals (Fig. 1B), in which cell proliferation is limited.

Moreover, while tardigrade embryos within the same exuvia typically develop synchronously (*25*), exposure to 10 µg/mL zeocin disrupted this coordination. Embryos from the same clutch displayed marked variation in developmental stage at a given timepoint (Fig. S11). To quantify this asynchrony, we incubated gravid adults about to lay in 10 µg/mL zeocin, fixed the embryos after three days, stained their nuclei, and imaged them by confocal microscopy. Based on nuclear number and distribution, we found variable degrees of asynchrony within over 90% of exuviae analyzed (Fig. 5D, E). In parallel, we tracked the long-term outcome of these embryos: all that eventually hatched under these conditions matured into fertile adults that fed normally, produced viable offspring, and maintained reproductive competence across generations (Fig. S12). These results show that, while tardigrade embryos exposed to sustained DNA damage can tolerate significant developmental delays, likely relying on active DNA repair, they nonetheless display signs of harm and can only withstand modest doses relative to adults.

We hypothesized that the differences in zeocin sensitivity manifested by highly replicative embryos and poorly proliferating adults would be underpinned by distinct DNA repair transcriptional programs. To test this, we extracted RNA from embryos and adults treated with the non-lethal dose of 10 µg/mL zeocin for three days then performed RT-qPCR on a subset of genes spanning five major DNA repair pathways (Fig. 5F, G). In agreement with prior results at a higher dose and shorter exposure (1 mg/mL, 24 hours; Fig. 1D), adults preferentially upregulated *PolQ*, *XRCC5*, and *Lig1*, consistent with reliance on NHEJ and BER (Fig. 5F). In contrast, embryos significantly upregulated *MSH6*, *ATR*, *Rad51*, *PolQ*, and *Lig1*, with the strongest responses observed in the mismatch repair (MMR) and HDR pathways (Fig. 5G). *PolQ* was upregulated in both embryos and adults, suggesting activation of the MMEJ pathway and/or a role in repairing DNA DSBs during mitosis (*28*, *29*). The strong upregulation of *MSH6*, *ATR* and *Rad51* in embryos points to a devoted, replication-coupled repair program (*30*, *31*).

Altogether, these findings demonstrate that embryos, while highly resilient, are particularly vulnerable to genotoxic stress, likely due to their elevated levels of replication. In response to DNA damage, they activate a distinct, replication-associated set of repair genes.

More broadly, our data support the view that replication constitutes a key window of vulnerability that limits the otherwise remarkable DNA damage resistance of tardigrades.

### Cross-species analyses reveal that proliferation levels modulate mortality kinetics under DNA damage

To ask if replication-associated vulnerability to DNA damage is conserved across animal lineages, we performed survival assays in two phylogenetically distant species: the nematode *Caenorhabditis elegans* and the planarian *Schmidtea mediterranea* (Fig. S13A–B). These species differ markedly in adult proliferative capacity: *C. elegans* is eutelic, lacking somatic cell division in adulthood (*32*, *33*), whereas *S. mediterranea* harbors neoblasts—adult pluripotent stem cells that divide continuously to maintain tissue homeostasis (*34*).

Exposing adults of both species to 1 mg/mL zeocin revealed strikingly different survival kinetics (Fig. S13C). *C. elegans*, which is much more short-lived under untreated conditions, showed an early decline, reaching complete lethality by day 8, while *S. mediterranea* remained viable until day 14 before a rapid collapse in survival between days 14 and 18. Although the onset of mortality varied—likely reflecting species-specific traits—the pace of decline, whether gradual or abrupt, correlated with each organism’s replicative capacity. Indeed, when survival data, normalized to controls, were plotted relative to the onset of mortality, regression analyses revealed a clear correlation between death rates and proliferative activity (Fig. S13D): *S. mediterranea* exhibited a very steep decline, while eutelic *C. elegans* succumbed more slowly.

Consistent with this, *C. elegans* post-embryonic development, which requires several essential mitotic divisions (*32*), proved highly sensitive to zeocin: continuous exposure of adults and their progeny led to widespread larval arrest and death on zeocin plates in a dose-dependent manner (Fig. S13E). At just 10 µg/mL zeocin, nearly all larvae exhibited developmental arrest (Fig. S13E). Overall, these findings suggest that a major consequence of genotoxic stress is the depletion of proliferative capacity, ultimately leading to the breakdown of tissue homeostasis and driving organismal decline.

## Discussion

In this study, we probed why tardigrades—despite their impressive DNA repair capacity and extremotolerance—still succumb to genotoxic stress. Exposing *Hypsibius exemplaris* to the radiomimetic drug zeocin revealed that mortality is primarily driven by exhaustion of gut cell replication capacity. Continuous zeocin treatment impairs gut cell replication, preventing their renewal and ultimately leading to midgut failure and death. Pulse–wash experiments (Fig. 4) further confirmed that even transient DNA damage in dividing gut cells is irreversible, resulting in collapse of tissue homeostasis in the mid-to-long term. Notably, this vulnerability associated with compromised DNA replication persists even as DNA damage triggers a strong, systemic induction of repair genes (*8–11*). These findings point to DNA replication, not repair, as a central bottleneck for survival under genotoxic challenge.

### The obligate passage through replication renders gut cells a major vulnerability

While our data cannot distinguish whether cells are most vulnerable to DNA damage during active DNA replication or because damaged cells subsequently lose their ability to replicate, it is clear that the incapacity to accomplish the obligatory transit through S-phase renders cycling cells a weakness in tardigrades.

It has been shown that, under starvation, *H. exemplaris* gut epithelial cells continue to proliferate, fully replacing the midgut lining within 5–6 days and thus preserving gut integrity (*13*). This contrasts with the response of storage cells—the other major tardigrade somatic replicative population—which, as we previously demonstrated, entirely cease replication events during food deprivation (*12*). This nutritional setup already emphasized the indispensable role of gut cell replication. Here, we show that zeocin exposure halts gut cell replication, and in agreement with the notion that their renewal is essential, this comes at a high price. Irreversible damage to gut cells following IR had already been postulated in tardigrades (*8*), but the possibility that this underlies animal death had not been further explored. Our results show that damaged cells fail to replicate and thus are not replaced, presumably disrupting digestion, ion balance, microbial homeostasis, and nutrient absorption. These effects likely contribute to the animals’ decline, especially in the context of systemic DNA damage, as DNA repair itself is highly energy-consuming (*35*, *36*).

We previously reported that storage cells replicate only during molting in both juveniles and adults, though this activity is more pronounced in juveniles (*12*). Importantly, preventing this replication using the deoxyribonucleotide reductase inhibitor hydroxyurea causes mortality during molting (*12*). Hydroxyurea acts by depriving already S-phase-committed cells of deoxyribonucleotides, stalling progressing replication forks (*37*). This further reinforces the idea that DNA replication represents a key vulnerability in tardigrades. In the present study, we found that zeocin markedly reduces the number of EdU⁺ storage cells (Fig. 3D–E), yet juveniles do not exhibit greater vulnerability to zeocin than adults (Fig. S7). Together, these observations raise the possibility that storage cells do not engage in replication at all during zeocin treatment, reinforcing our conclusion that mortality stems primarily from the loss of gut cell replicative activity. We hypothesize that storage cells, likely arrested in the G_1_ or G_0_ phase of the cell cycle between molting episodes, simply do not receive—or ignore—the signal to proceed into S phase, thereby avoiding engagement in DNA replication. This would support the view that DNA damage in tardigrades is not inherently lethal unless replication is actively attempted.

We wish to emphasize, however, that beyond DNA breaks, zeocin-induced oxidative damage (*16*) may also affect proteins and lipids, and its genotoxicity extends beyond the gut. For instance, we observed nuclear fragmentation and disrupted cuticle architecture in epidermal tissues (Fig. 2D; Fig. S6). These systemic effects suggest that mortality during constant zeocin treatment is not solely due to midgut dysfunction but probably reflects an overall collapse of organismal homeostasis in the face of accumulating lesions.

### Progenitor cells and tissue homeostasis

Our spatial EdU labeling data (Fig. 4) support the existence of dedicated gut progenitor cells located at the anterior and posterior edges of the midgut (*13*, *24*). These putative unipotent stem cells likely sustain continuous midgut epithelial renewal. Their progressive loss—or complete loss of replication capacity—after zeocin exposure suggests that they are particularly vulnerable to genotoxic stress. This observation aligns with findings from other systems, such as mammalian adult intestinal stem cells, which are key drivers of intestinal epithelial homeostasis (*38*). In tardigrades, as in mammals, the loss of replicative progenitors may similarly underlie progressive tissue degeneration and organismal decline.

The depletion of these putative gut progenitor cells parallels what we observed in germ cells: here too, DNA damage dose- and time-dependently impaired fertility (Fig. 5A–B; Fig. S10). This reinforces the idea that stem-like cells, whether somatic or germline, are particularly sensitive to DNA damage due to their proliferative nature and their need to maintain genomic integrity for tissue renewal and reproductive success (*39*, *40*).

Animal embryonic development and larval stages are also highly proliferative, setting the stage for differentiation into adult tissues. As such, these stages are expected to be more vulnerable to DNA damage. We confirmed this in *H. exemplaris* embryos (Fig. 5C–E) and in *C. elegans* larvae (Fig. S13E). In *S. mediterranea* adults, which rely on pluripotent stem cells called neoblasts for tissue renewal (*34*), zeocin-induced DNA damage triggered a sharp loss of viability, with animals dying abruptly after two weeks of treatment (Fig. S13C). This phenotype likely reflects neoblast depletion, since IR is known to ablate neoblasts and drive *S. mediterranea* death within three weeks (*41*, *42*). By contrast, adult *C. elegans*, with their eutelic somatic tissues, exhibit a more gradual survival decline under DNA damage than *S. mediterranea* (Fig. S13D). Together, these observations underscore that reliance on proliferative progenitors renders tissues acutely sensitive to genotoxic stress.

### Tissue- and stage-specific DNA repair responses in tardigrades

Recent studies have mapped the transcriptional landscape responding to DNA damage in *H. exemplaris* adults (*8*, *9*), and we confirm these findings here by profiling a subset of repair genes via RT-qPCR in response to zeocin (Fig. 1D). Importantly, we expand this knowledge by dissecting how distinct tissues and developmental stages respond to and are affected by DNA damage.

Within adults, EdU incorporation profiles revealed a heterogeneous landscape: while replication ceased in basally proliferative cell types, some quiescent cells exhibited low-intensity EdU signals indicative of reparative DNA synthesis (Fig. 3; Fig. S8–S9). The distribution of these signals, together with the comparatively modest upregulation of *Rad51* relative to genes from other repair pathways (Fig. 1D), suggests that HDR is engaged in select subpopulations, including epidermal cells, cells in the brain and pharynx, and claw gland cells (Fig. 3B). HDR is typically restricted to S and G_2_ phases, as it requires a sister chromatid for accurate repair (*31*). This restriction is critical because HDR in G_1_ could erroneously use homologous—but non-identical—sequences, potentially leading to gross chromosomal rearrangements and loss of heterozygosity. Engagement of HDR is prevented in G_1_ by limiting DNA end resection, a step regulated by CDK-mediated phosphorylation of Sae2/CtIP (*43–45*).

Our observation that some tardigrade cells might engage HDR is intriguing because it suggests that quiescent cells may be arrested in G_2_ rather than residing in G_0_ or G_1_. While most non-proliferative cells are thought to remain in G_0_ after exiting G_1_ with a 2C DNA content (*46*), exceptions exist, usually related to polyploidy (*47–49*). Alternatively, HDR could engage atypically in G_1_. For instance, DNA breaks at the repetitive sequences conforming centromeres can recruit the HDR machinery in G_1_ through an H3K4me2-dependent pathway, protecting genome integrity but potentially triggering chromosomal rearrangements if misregulated (*50*). Whether G_2_ arrest or unconventional G_1_ HDR, the reparative EdU signals we detect highlight cell type-specific differences in quiescence states and repair profiles.

In embryos, a distinct DNA damage management strategy likely underpins their lower yet remarkable resilience. Despite developmental delays and asynchrony under continuous 10 µg/mL zeocin exposure, ∼80% of embryos still hatch and produce fertile progeny (Fig. 5C– E; Fig. S12). Unlike adults, which primarily rely on BER and NHEJ repair pathways, embryos preferentially activate MMR and HDR, suggesting a reliance on replication-coupled pathways to resolve DNA damage (*30*, *31*)—in line with their high mitotic activity. The observed upregulation of *POLQ*, encoding polymerase theta (Polθ), which has recently been shown to repair persistent DSBs during mitosis (*28*, *29*), contrasts with its classical designation as a backup pathway for NHEJ. This further supports the notion that embryos deploy DNA repair programs adapted to their intense proliferation.

### A universal link between DNA damage, fat loss and body size?

We reveal that a striking physiological output of DNA damage is a profound effect on body size, underlined by a dramatic fat loss (Fig. 2). In *C. elegans*, NER-deficient mutants exhibit growth defects and UV-induced metabolic adjustments that mimic starvation and aging, involving autophagy and AMPK signaling (*51*). Persistent DNA damage also induces mitochondrial β-oxidation and fat depletion (*52*). Mechanistically, we recently demonstrated in *S. cerevisiae* and human cells that repairing DNA breaks requires sterol sequestration into lipid droplets (*17*), thus potentially displacing triacylglycerols (TAG) from these stores. In agreement, recent evidence shows that TAG lipolysis is induced in response to DNA breaks (*53*). These processes may help understand why genome instability syndromes are associated to lipodystrophy in humans (*54*).

In this light, our findings in tardigrades suggest that body size is partly governed by the volume of adipose depots, particularly storage cells, whose lipid content sharply declines during nutrient deprivation and DNA damage stress (Fig. 2A–C). In previous work, we showed that juvenile *H. exemplaris* arrest growth under starvation (*12*). Here, we find that adults shrink under both starvation and zeocin exposure (Fig. 2A–B). Whether this shrinkage reflects a global decrease in cell size or is limited to specific cell types remains to be explored. Notably, storage cell size is markedly reduced in *H. exemplaris* under both starvation and zeocin treatment (Fig. 2A), as previously reported for starvation in other tardigrade species (*55*). However, alterations in total cell number—whether through cell loss or recycling—may also contribute.

Fat loss in tardigrades may occur early during zeocin treatment rather than being solely a consequence of feeding failure. Interestingly, we repeatedly observed zeocin-treated animals actively searching for food despite being unable to assimilate it, in marked contrast to starved animals, which enter an inactive state (*12*). This suggests that DNA damage might trigger a metabolic rewiring that mobilizes fat reserves independently of nutrient uptake. Future studies examining cellular dimensions, numbers, and fat depots under various conditions will be crucial to elucidate how genome instability and metabolic adaptations intersect to regulate body size and potentially influence stress resilience.

Finally, it is worth noting that while genetic instability is historically associated with deregulated cell proliferation and uncontrolled growth, as in cancer (*56*), growth retardation is also a well-documented feature of multiple genome instability syndromes in humans (*57*). Although body size reduction and growth retardation are not strictly analogous, our findings on zeocin-induced size reduction in *H. exemplaris* (Fig. 2A–B) align with this notion and open new avenues for exploring the links between genome instability and growth regulation.

## Conclusion

Our findings establish DNA replication as a critical Achilles’ heel in *H. exemplaris*, despite its extraordinary DNA repair arsenal. Proliferative gut cells, essential for midgut renewal, are irreversibly compromised by zeocin, leading to tissue failure and organismal death. Germ cells and embryos further underscore replication-associated vulnerability: germ-cell DNA damage abolishes fertility, while highly proliferative embryos are more sensitive to genotoxic stress than adults despite engaging distinct, replication-focused repair programs. These insights into replication-linked sensitivity in an extremotolerant organism may inform future strategies in regenerative medicine, aging, and cancer biology, where targeting proliferative activity remains a key therapeutic avenue. In sum, tardigrades illustrate that survival under genotoxic stress is not only about resisting and repairing DNA damage, but also about preserving the potential to safely replicate DNA and thus maintain tissue homeostasis.

## Material and Methods

### *Hypsibius exemplaris* husbandry and zeocin treatments

*H. exemplaris* (Z151 strain) was maintained as previously described (*12*). Briefly, tardigrades were cultured in an incubator at 15 °C in 55 mm plastic Petri dishes filled halfway with spring water (Volvic) filtered through a 0.2 µm mesh. To promote movement, the dish bottoms were lightly scratched with sandpaper. Animals were kept under a 12 h light/12 h dark photoperiod and fed *ad libitum* with *Chlorococcum sp*. algae, which were grown in 50 mL tubes using BG-11 growth medium (Gibco, A1379901). Water changes were performed every 2–3 weeks to maintain optimal conditions.

All experimental incubations lasting ≥24 h were conducted under identical conditions in 35 mm Petri dishes and placed inside cardboard boxes to protect photosensitive compounds from light degradation. Juvenile and adult tardigrades, as well as embryos, were incubated with the radiomimetic agent zeocin (R25001, ThermoFisher) at the specified concentrations. For long-term continuous zeocin exposure, the drug was renewed every 7 days to ensure sustained activity, as confirmed by monitoring its efficacy in a heterologous yeast system (Fig. S2). Pulse treatments were applied for the indicated durations, followed by thorough washing with filtered spring water (FSW) to remove residual zeocin.

#### Survival, fertility, and hatchability assessment

For all experiments, animals were selected at early to late vitellogenesis stages, as defined by (*58*). Adults measured ≥240 µm in length, while juveniles ranged between 120– 180 µm.

Survival was assessed by calculating the percentage of animals alive at each timepoint under a given zeocin condition, following previously established criteria (*2*).

Fertility was measured as the number of eggs laid per animal over a defined period. For each timepoint, fertility was calculated by dividing the total number of eggs laid by the number of live animals at that time (using the last recorded number of live animals when all animals had died). In Fig. 5A–B, fertility is presented as the mean cumulative number of eggs laid per animal over time; in Fig. S9, it is shown as the mean number of eggs laid per animal at each timepoint.

Hatchability was defined as the proportion of embryos successfully completing development and hatching from their eggshells. It was calculated by dividing the number of hatched eggshells at each timepoint by the total number of eggs at the beginning of the experiment. To ensure continuous zeocin exposure throughout development, gravid animals about to molt were used. As *H. exemplaris* is parthenogenetic, embryos initiate development immediately after oviposition without requiring fertilization.

#### DAPI, BODIPY, and MemBrite staining

Animals were collected, filtered through a 40 µm mesh, and transferred to a glass depression slide within a humid chamber using a glass Pasteur pipette. Whenever possible, samples were kept in the dark throughout the procedure to protect light-sensitive reagents.

For neutral lipid labeling, live specimens were incubated for 30 min in 10 µg/mL BODIPY (Difluoro{2-[1-(3,5-dimethyl-2H-pyrrol-2-ylidene-N)ethyl]-3,5-dimethyl-1H-pyrrolato- N}boron; Sigma-Aldrich, 790389) diluted in FSW. Following incubation, animals were rinsed three times in FSW and fixed for 1 h at room temperature (RT) in 4% paraformaldehyde (PFA) in 1x PBS. Fixed samples were washed three times for 15 min each in 1x PBS, then incubated for 30 min in 1 µg/mL DAPI (4′,6-diamidino-2-phenylindole; Sigma-Aldrich, D9542) diluted in 1x PBS to stain nuclei. After DAPI staining, samples were washed three additional times in 1x PBS (15 min per wash), mounted on slides with ProLong™ Gold Antifade Mountant (Invitrogen, P36930), and imaged by confocal microscopy.

Cuticular structures were stained using the MemBrite™ Fix 543/560 Cell Surface Staining Kit (Biotium, BTM30094). Fresh pre-staining and staining solutions were prepared at 2.5x concentration from 1000x stock in 200 µL FSW. Animals were first incubated in pre-staining working solution for 10 min, then transferred to the staining working solution for 15 min. After staining, animals were washed three times in FSW, fixed for 1 h in 4% PFA in 1x PBS, washed in 1x PBS as above, and mounted in ProLong™ Gold Antifade Mountant for imaging. All incubations were performed at RT.

#### EdU experiments

To detect replicative and reparative DNA synthesis, animals were incubated in 100 µM EdU (Click-iT™ EdU Cell Proliferation Kit for Imaging, Alexa Fluor™ 488 Dye; Invitrogen, C10337) diluted in FSW containing algae for experiment-specific durations. Following EdU exposure, tardigrades were collected, transferred to a glass depression slide, and anesthetized with 10% ethanol for 15 min to prevent contraction during fixation. Animals were then fixed for 1 h at RT in 4% PFA in 1x PBS supplemented with 1% Triton X-100 (PTx).

After fixation, samples were washed three times for 15 min each in PTx and blocked for 2 h in 0.2 µm-filtered blocking solution containing 10% bovine serum albumin (BSA) in 1x PBS. The Click-iT™ EdU detection reaction was performed for 1 h at RT according to the manufacturer’s protocol. Following detection, samples were washed four times for 10 min each in 1x PBS + 0.1% Tween-20 (PTw). Nuclei were counterstained with DAPI (1 µg/mL), and specimens were mounted in ProLong™ Gold Antifade Mountant (Invitrogen, P36930) for confocal imaging, as described above.

#### Comet assay for tardigrade storage cells

The alkaline comet assay was performed as previously described (*59*), with adaptations from a protocol established for tardigrade storage cells (*20*). A detailed experimental procedure is provided in Supplementary File S1. Fluorescence images were acquired on a Leica Thunder microscope (20x objective) with identical exposure settings across all samples. Comet tail length and percent DNA in the tail were quantified using CometScore 2.0. While control slides consistently displayed some levels of tailing, likely reflecting intrinsic assay background, zeocin-treated samples showed a significant and reproducible increase in both parameters, confirming an increase in DNA strand breaks.

#### RT-qPCR

Total RNA was extracted from adults and embryos using the RNAqueous-Micro Total RNA Isolation Kit (Ambion, AM1931), according to the manufacturer’s instructions. For each condition in every experiment, 150–250 adults or 250–400 embryos were collected for RNA extraction. Genomic DNA was removed by DNase treatment following the manufacturer’s recommendations. Reverse transcription was performed using SuperScript™ II Reverse Transcriptase (Invitrogen, 18064014) with random primers. Quantitative PCR was carried out using Platinum Taq Polymerase (Invitrogen, 10966-034) and a home-made SYBR Green qPCR master mix (*60*). Reactions were run on a Stratagene Mx3000P system (Agilent).

For each gene and condition, transcript levels were quantified from three technical replicates and normalized to the housekeeping gene *Ef1alpha*. Relative expression was calculated using the ΔΔCt method, comparing experimental conditions to untreated controls. Data are reported as arbitrary units (A.U.) in all figures. Three independent biological replicates were performed for each experiment. Primer sequences are provided in Supplementary File S2.

#### Imaging, length measurements, cell counting, and asynchrony assessment

Live tardigrades were anesthetized in 20 mM levamisole hydrochloride (MedChemExpress, HY-13666) in Milli-Q H₂O, mounted on glass slides, and covered with coverslips supported at the corners with clay to prevent compression. Differential interference contrast (DIC) color images were acquired using a Leica K3C digital camera mounted on a Leica Thunder light microscope. Body length measurements were performed in ImageJ (*61*), from the head to the juncture at the posterior-most segment bearing legs (*12*). For starved tardigrades, measurements were taken immediately after they came back to an active state to ensure full body extension. Live images in Fig. S4, S5, and S12 were captured directly in culture medium using a smartphone camera mounted on a stereoscope.

Fluorescence images were acquired on a Zeiss LSM980 confocal microscope equipped with an Airyscan2 module. Identical scanning parameters were applied across conditions within each experiment for consistency, except for Fig. 3B and Fig. S8, where laser power and gain were increased to visualize the weaker signals of reparative DNA synthesis. Confocal Z-stacks were processed and brightness/contrast adjusted in ImageJ using K. Terretaz’s visualization toolset (https://github.com/kwolbachia/Visualization_toolset).

EdU fluorescence quantification (Fig. S9) was performed by manually outlining nuclei using DAPI counterstaining and measuring mean gray values in the EdU channel in ImageJ. EdU⁺ cells were manually counted from full Z-stacks spanning the entire animal depth.

For embryo asynchrony assessment, only exuviae containing ≥3 embryos were analyzed. After three days of development under control or 10 µg/mL zeocin conditions, fixed and DAPI-stained embryos within their exuviae were imaged and classified into three categories based on approximate nuclear number and distribution: synchronous, one asynchronous embryo, or ≥2 asynchronous embryos. Tardigrade schematics and figure layouts were created in Adobe Illustrator.

#### Experiments with heterologous systems

Yeast culture, protein extraction, and western blotting were performed as previously described (*17*).

*C. elegans* worm N2 strain (wildtype ancestral, Bristol) was obtained from the CGC (https://cgc.umn.edu) and maintained at 20 °C on NGM plates seeded with *E. coli* HT115, following standard procedures (*62*). For zeocin treatment, the drug was incorporated into both the agar plates and the bacterial lawn. Plates and bacterial cultures were buffered at pH 7.4 with 20 mM HEPES to preserve zeocin activity. To prevent zeocin degradation, *E. coli* were heat-inactivated by concentrating saturated HT115 cultures 30-fold and incubating for 30 min at 65 °C. This bacterial suspension was then used to seed NGM plates with or without zeocin. Worms were synchronized using the alkaline bleach method (1.2% NaOCl, 250 mM KOH in water), which preserves only eggs protected by their eggshells. Eggs were allowed to hatch overnight at 16 °C in M9 buffer (3 g KH₂PO₄, 6 g Na₂HPO₄, 5 g NaCl, 1 mM MgSO₄ per liter H₂O) in the absence of food. Synchronized larvae were then transferred to seeded plates and allowed to reach adulthood (∼64 h at 19 °C) before being transferred onto zeocin or control plates. Survival was monitored daily under a dissecting scope, and worms were transferred to fresh plates every three days to maintain culture conditions and constant zeocin levels. To evaluate the effect of zeocin treatment on *C. elegans* exhibiting somatic cell proliferation, the progeny of adult worms treated for 24 h with various zeocin concentrations was analyzed. Among the eggs that hatched, we scored the number of larvae that reached adulthood after 72h in the presence or absence of zeocin at different concentrations.

Intact, homeostatic planarians from an asexual clonal line of the species *Schmidtea mediterranea* were continuously exposed to 1 mg/mL zeocin diluted in planarian artificial water (*63*) and kept at 20°C in the dark. Viability was scored every other day under a stereomicroscope. Three biological replicates were performed, each comprising 20 planarians.

#### Statistics

The normality of all datasets was assessed using the Shapiro–Wilk test. For all two-group comparisons, the nonparametric Mann–Whitney U test was applied. For RT-qPCR analyses, an ordinary one-way ANOVA on ΔΔCt values with Dunnett’s multiple comparisons test was performed. For all other multiple group (>2) comparisons, the Kruskal–Wallis test followed by Dunn’s multiple comparisons test was used. Statistical significance is represented as follows: *p* ≤ 0.05 (*), *p* ≤ 0.01 (**), *p* ≤ 0.001 (***), and *p* ≤ 0.0001 (****). All graphs and statistical analyses were generated using GraphPad Prism 10.

## Acknowledgements

We thank Kseniya Samardak for her help with tardigrade husbandry and Sylvain Kumanski for having performed the western blots of Rad53. We are grateful to Kevin Terretaz for his valuable tips and macros on ImageJ. We acknowledge the imaging facility MRI, member of the France-BioImaging national infrastructure supported by the French National Research Agency (ANR- 10-INBS-04, «Investments for the future»), and we thank the program Montpellier Université d’Excellence (MUSE I-SITE) as well as the Program Impact Santé France 2030 for funding this research.

## Authors’ contributions

Conceptualization: G.Q.-A., M.M.-C.; Methodology: G.Q.-A., B.L., MD.M.; M.M.-C.; Validation: G.Q.-A., P.F., B.L., MD.M.; Formal analysis: G.Q.-A.; Investigation: G.Q.-A., P.F., B.L., MD.M.; Writing - original draft: G.Q.-A., M.M.-C.; Writing - review & editing: G.Q.-A., M.M.-C., B.L., MD.M.; Visualization: G.Q.-A.; Supervision: G.Q.-A., M.M.-C.; Project administration: G.Q.-A., M.M.-C.; Funding acquisition: M.M.-C.

## Data availability

All data are available within the paper and supplementary material.

## Supplementary Figures

**Figure S1.**
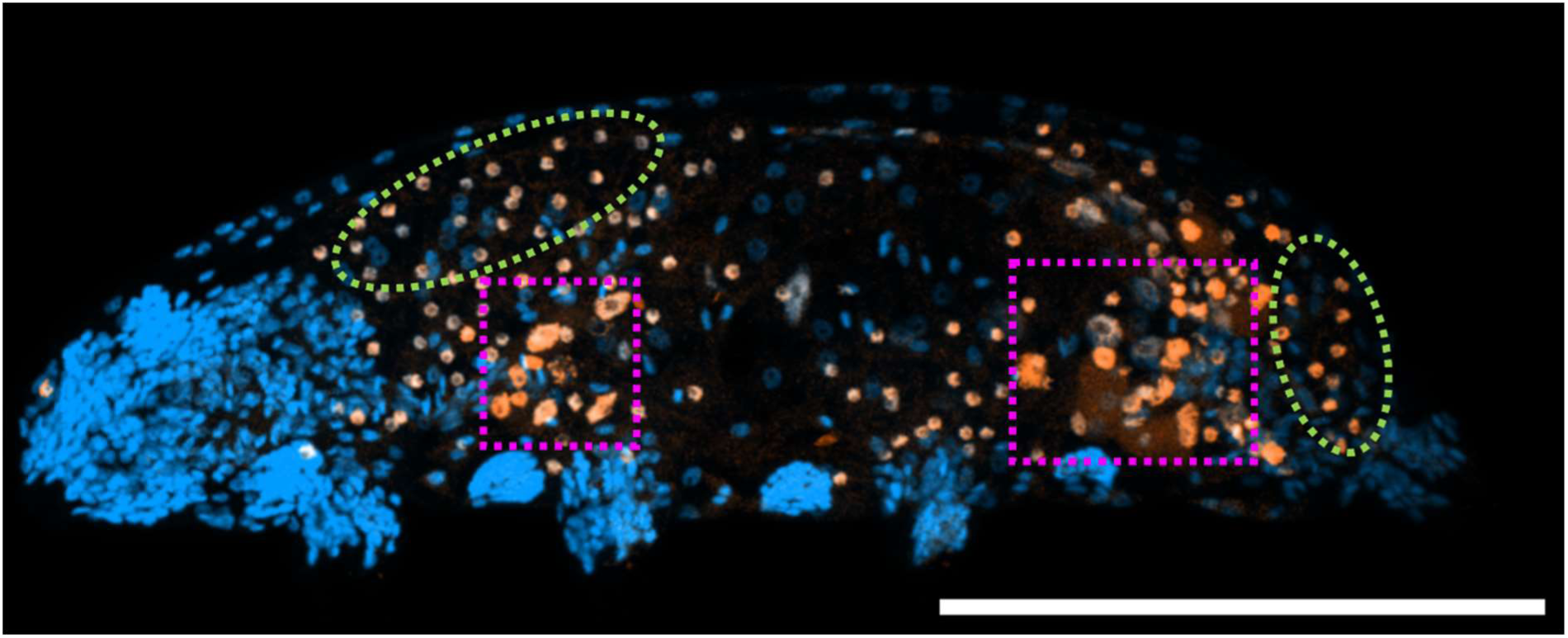
Two-week EdU incubation reveals somatic proliferative cell types in tardigrades. Representative maximum projection image of a tardigrade following a two-week EdU exposure. Nuclei are labeled with DAPI (blue), and proliferative (EdU⁺) cells are shown in orange. Green circles highlight EdU⁺ storage cells located in both anterior and posterior regions, while magenta squares indicate EdU⁺ cells in the foregut and hindgut. Scale bar = 100 μm.

**Figure S2.**
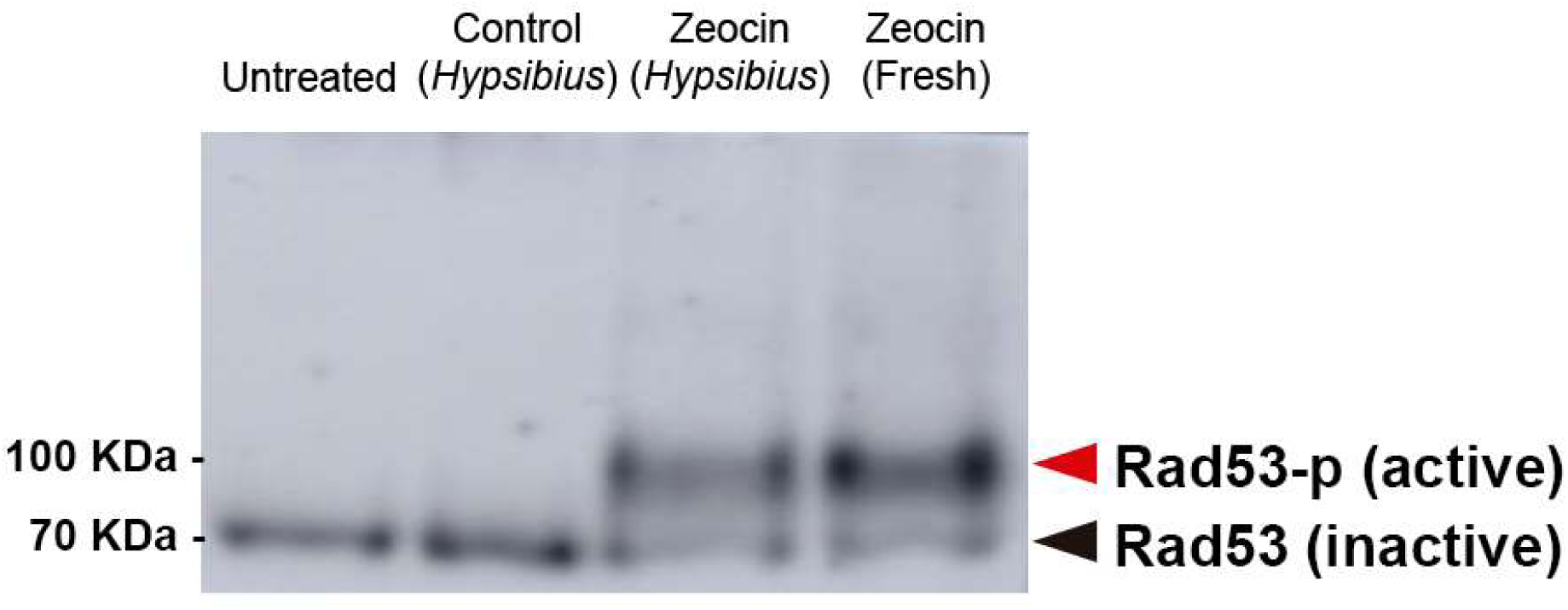
Zeocin retains genotoxic activity in tardigrade culture medium for at least seven days. Western blot of phosphorylated Rad53 (Rad53-p) in *S. cerevisiae* protein extracts. Black arrow represents Rad53 and red arrow represents Rad53-p, indicating an activation of the DNA Damage Response (DDR). Untreated = untreated yeast. Control (*Hypsibius*) = yeast treated with a 1/100 dilution of the water in which untreated *H. exemplaris* were incubated for one week. Zeocin (*Hypsibius*) = yeast treated with a 1/100 dilution of the 1 mg/mL condition in which *H. exemplaris* were incubated for a week [final concentration = 10 μg/mL]. Zeocin (Fresh) = yeast treated with fresh zeocin at 10 μg/mL. All yeast treatments were carried out for 3 hours prior to protein extraction.

**Figure S3.**
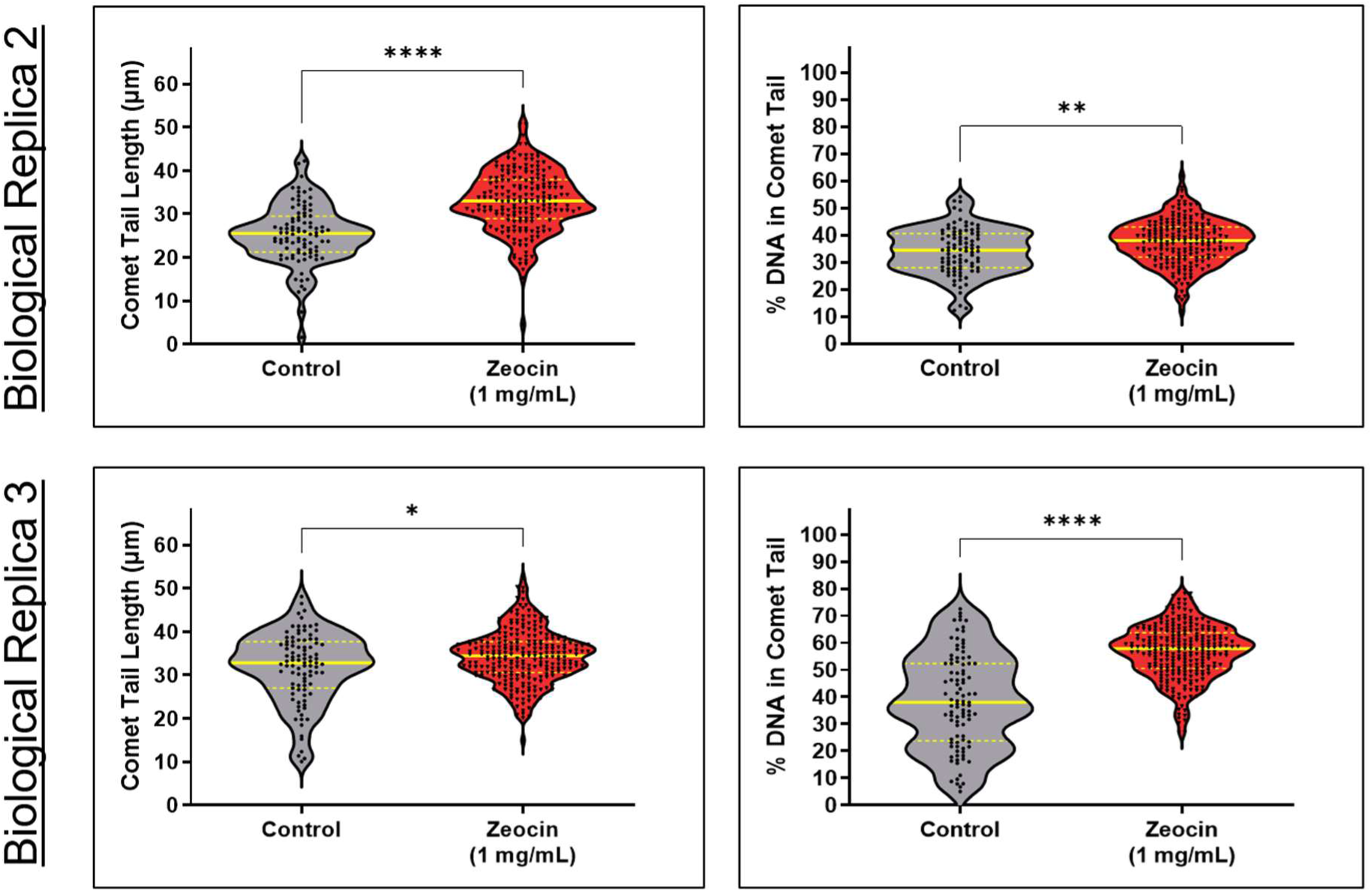
Zeocin exposure induces DNA damage in tardigrade storage cells. Violin plots showing comet tail length and percentage of DNA in the comet tail, comparing control and zeocin-treated (1 mg/mL for 4 days) storage cells from two biological replicates. In replicate 2, *n* = 92 (control) and *n* = 192 (zeocin-treated); in replicate 3, *n* = 95 (control) and *n* = 255 (zeocin-treated). Plot layout and quantification follow the format of Fig. 1F–G. **** *p* ≤ 0.0001; ** *p* ≤ 0.01; * *p* ≤ 0.05 (Mann–Whitney U test).

**Figure S4.**
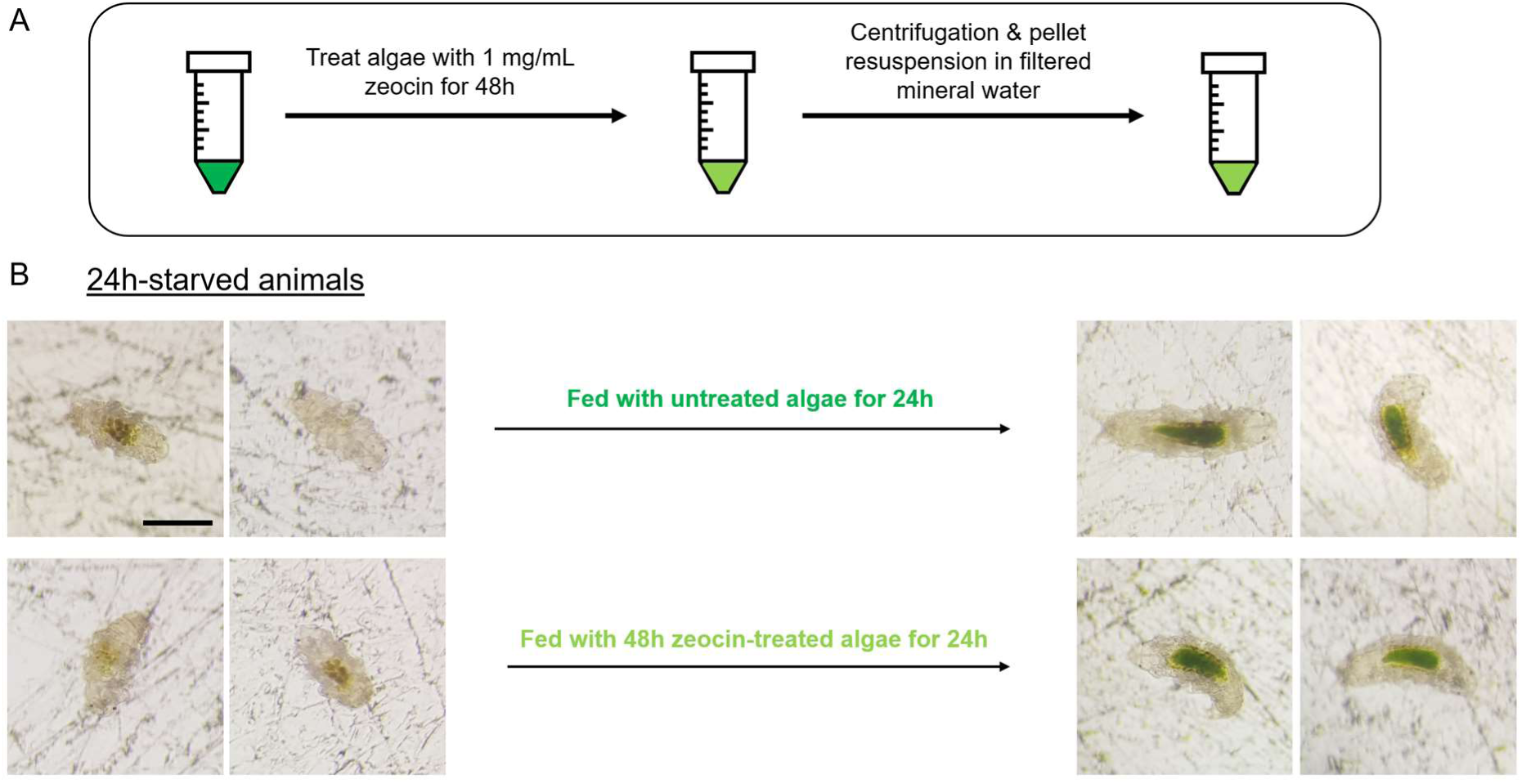
Untreated H. exemplaris consume zeocin-treated algae. (**A)** Schematic illustrating the treatment of *Chlorococcum* sp. algae with 1 mg/mL zeocin for 48 h, followed by centrifugation and resuspension in filtered mineral water for feeding to untreated tardigrades. The light green color denotes zeocin-treated algae, which show visible color decay after treatment. (**B)** Representative images of *H. exemplaris* specimens starved for 24 hours (left) and fed with either untreated or zeocin-treated algae (right). In both cases, the animals’ guts are visibly filled with algae, indicating that untreated tardigrades are capable of ingesting zeocin-treated algae. *n* > 30 in all cases. Scale bar = 100 μm.

**Figure S5.**
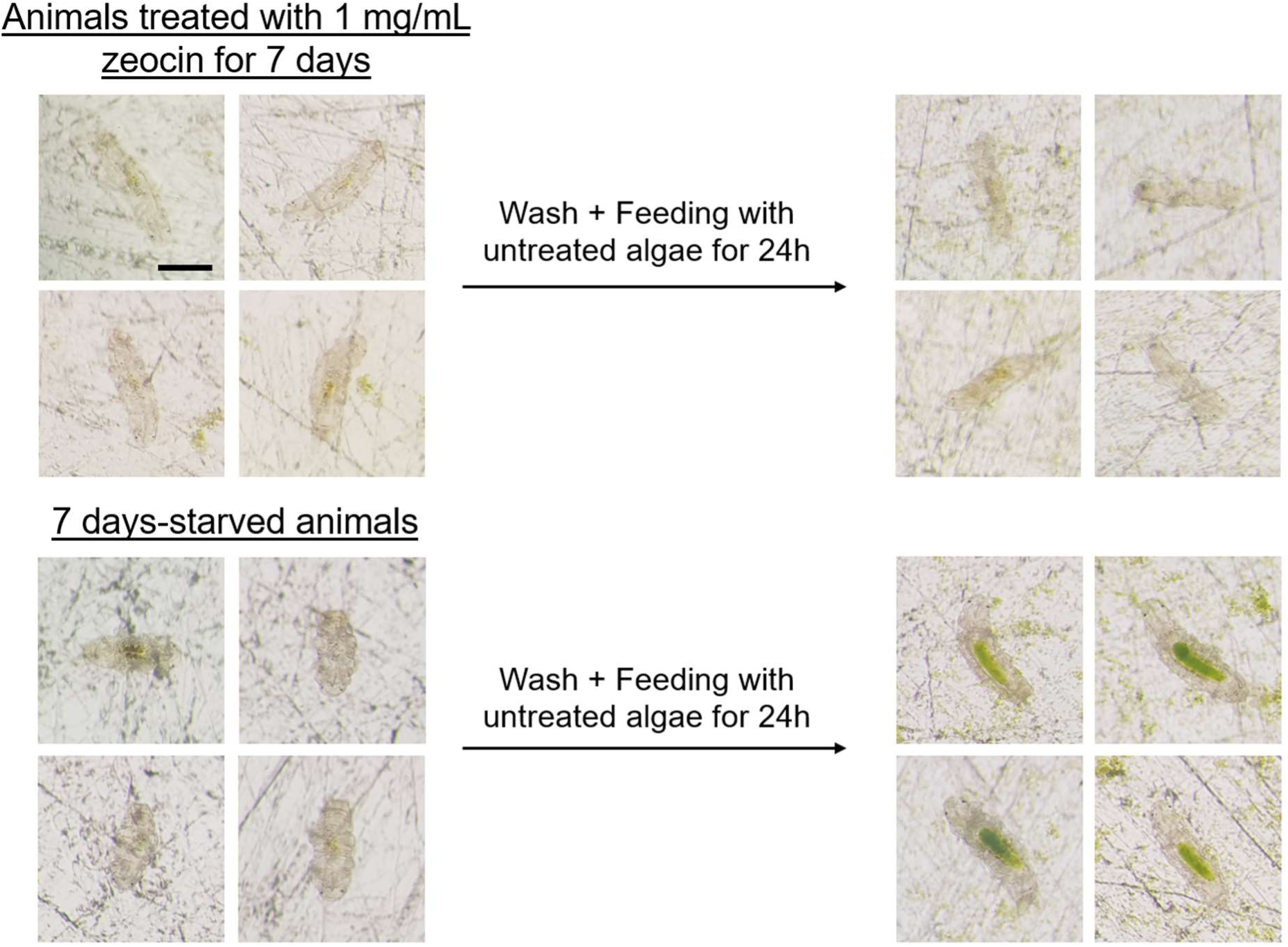
Extended zeocin exposure impairs feeding in H. exemplaris. Representative images of *H. exemplaris* specimens treated with 1 mg/mL zeocin for 7 days (top left), followed by 24 hours of feeding with untreated *Chlorococcum* sp. algae (top right). These animals show no visible algae in their guts, indicating a loss of feeding ability. In contrast, animals starved for 7 days without zeocin exposure (bottom left), then fed algae for 24 hours (bottom right), display guts filled with green algae, demonstrating preserved feeding capacity. *n* > 30 in all cases. Scale bar = 100 μm.

**Figure S6.**
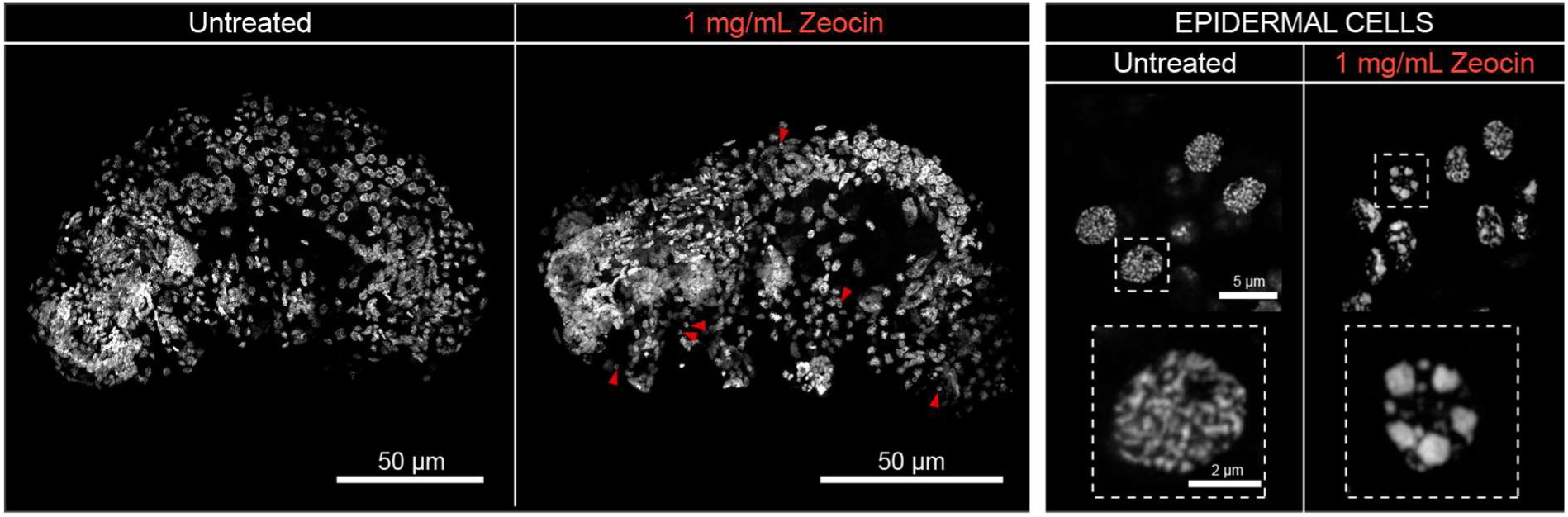
Effects of zeocin on the DNA of *H. exemplaris* epidermal cells. Left: Maximum projection images of whole animals stained with DAPI (nuclei, white) following a 1-week exposure to either untreated or 1 mg/mL zeocin conditions. Red arrowheads indicate affected nuclei in epidermal cells. Right: High-magnification images of epidermal cell nuclei from adult *H. exemplaris* under the same conditions, highlighting DNA fragmentation in zeocin-treated samples. Scale bars as indicated.

**Figure S7.**
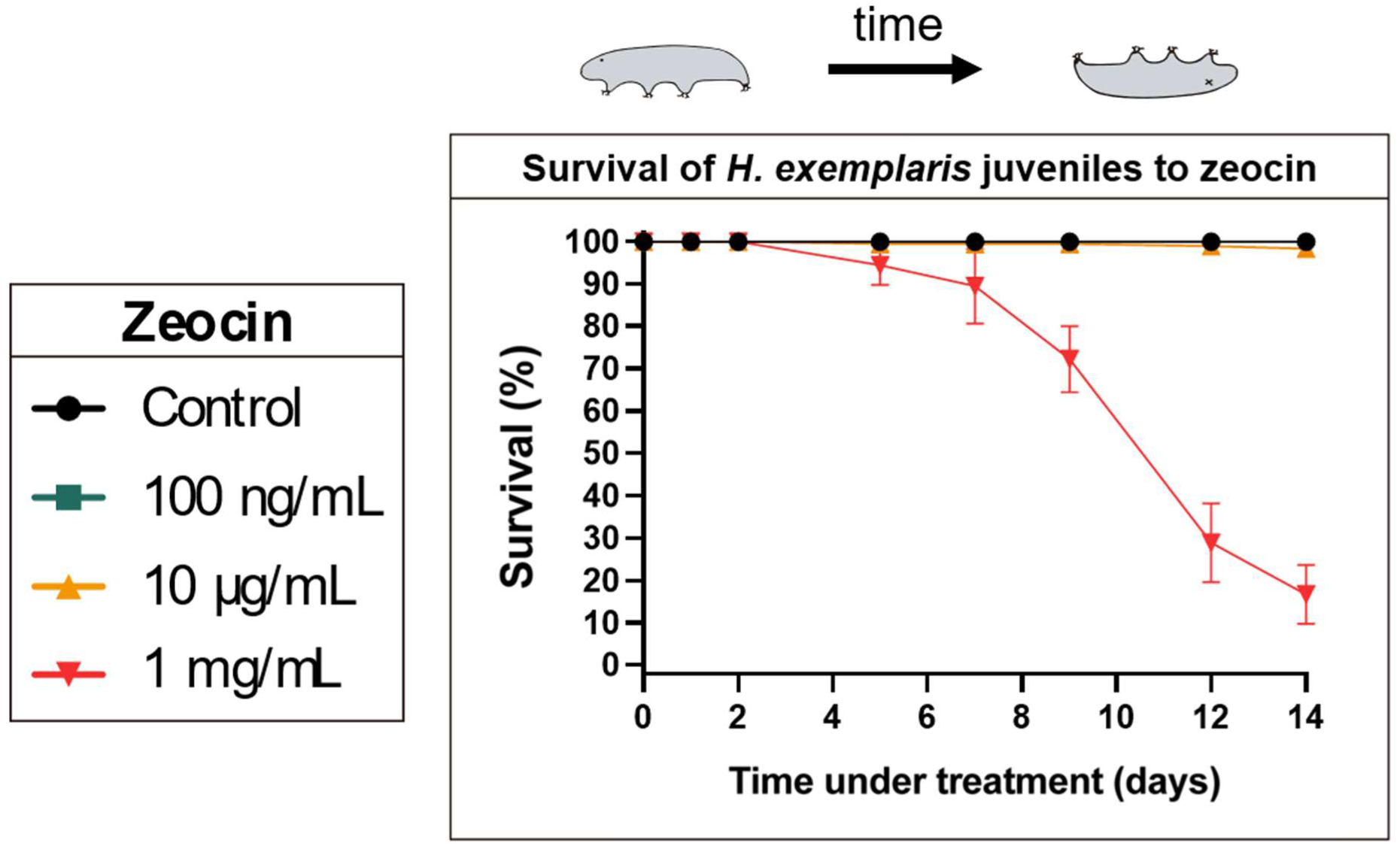
Juveniles of *H. exemplaris* exhibit similar sensitivity to zeocin as adults. Survival curves of *H. exemplaris* juveniles over time under continuous zeocin treatment at varying doses. Data are presented as the mean ± standard error of the mean (SEM) from three biological replicates, each consisting of 60 animals.

**Figure S8.**
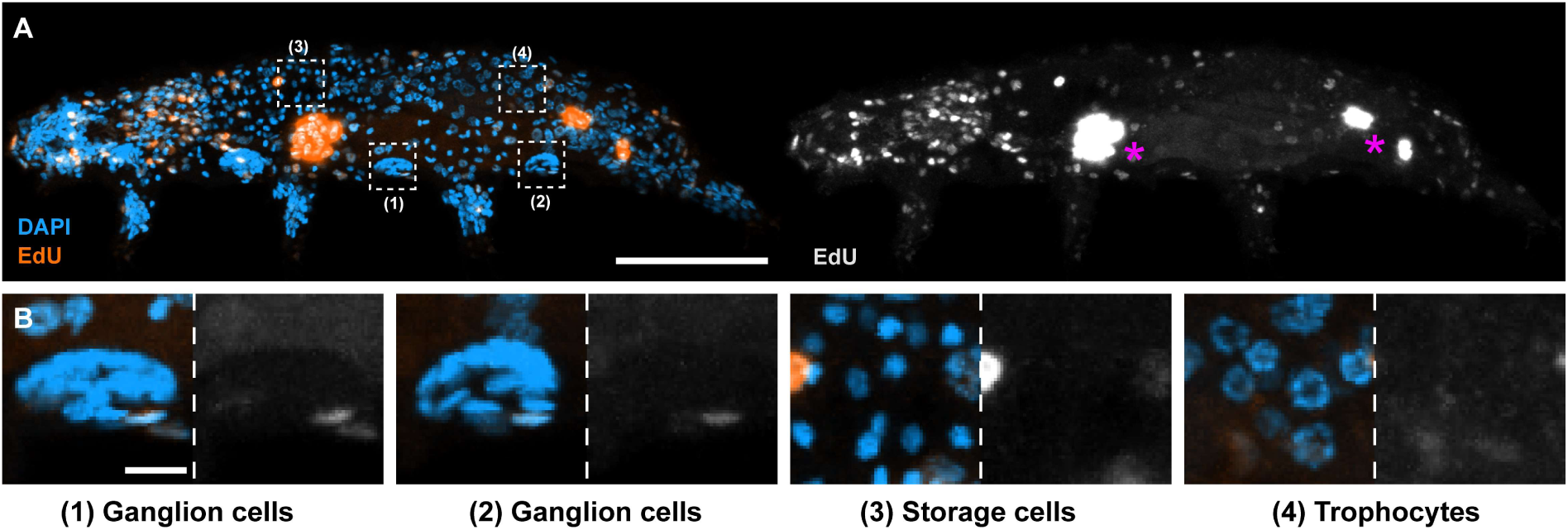
Zeocin induces reparative DNA synthesis in select, but not all, cell types. (A) Representative images of *H. exemplaris* after 4-day EdU + zeocin treatment, showing nuclei (DAPI, blue) and EdU^+^ cells (orange). Right panel displays EdU signal alone (gray). Magenta asterisks mark EdU^+^ gut cells in the foregut and hindgut. Dashed boxes highlight regions shown in (B). (B) Enlarged views of specific cell types from the boxed areas in (A). Note the absence of EdU incorporation in these cell types. Images are maximum intensity projections of full-depth z-stacks. Scale bars: 50 μm (A); 5 μm (B).

**Figure S9.**
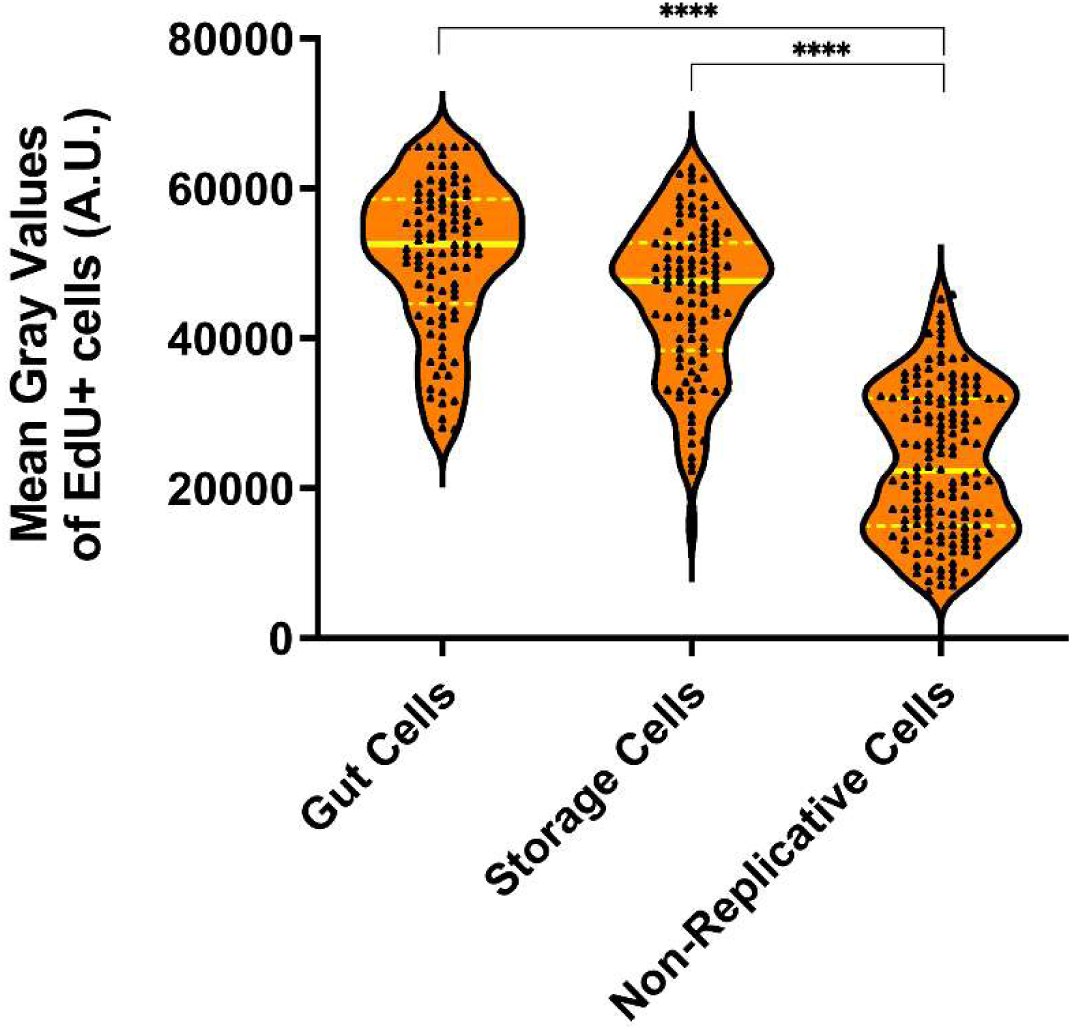
EdU^+^ non-replicative cells display lower fluorescence levels. Violin plots showing EdU fluorescence intensity in gut and storage cells (replicative), and in other somatic cell types (non-replicative), following zeocin exposure (as in Fig. 3A, C). Approximately 10 cells were quantified per animal from at least three animals across three independent experiments. *n* = 100 (gut cells), *n* = 101 (storage cells), and *n* = 150 (non-replicative cells). Plot layout and quantification follow the format used in Fig. 1F–G. Each cell is represented by a triangle. A.U. = arbitrary units. **** *p* ≤ 0.0001 (Kruskal–Wallis test).

**Figure S10.**
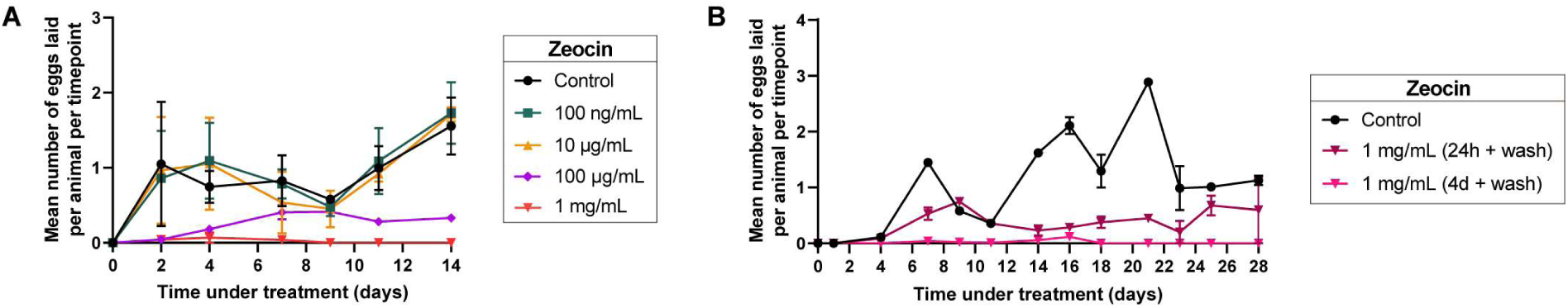
Fertility of zeocin-treated *H. exemplaris* measured per timepoint. (A) Average number of eggs laid per alive animal per timepoint over 14 days of continuous zeocin treatment at varying doses. Data are shown as mean ± standard error of the mean (SEM) from three biological replicates, each consisting of 60–120 animals. (B) Average number of eggs laid per alive animal per timepoint over 28 days following two pulsed zeocin treatment regimens (24 h pulse + wash; 4-day pulse + wash) at 1 mg/mL. Data are shown as mean ± SEM from two biological replicates, each consisting of 50 animals.

**Figure S11.**
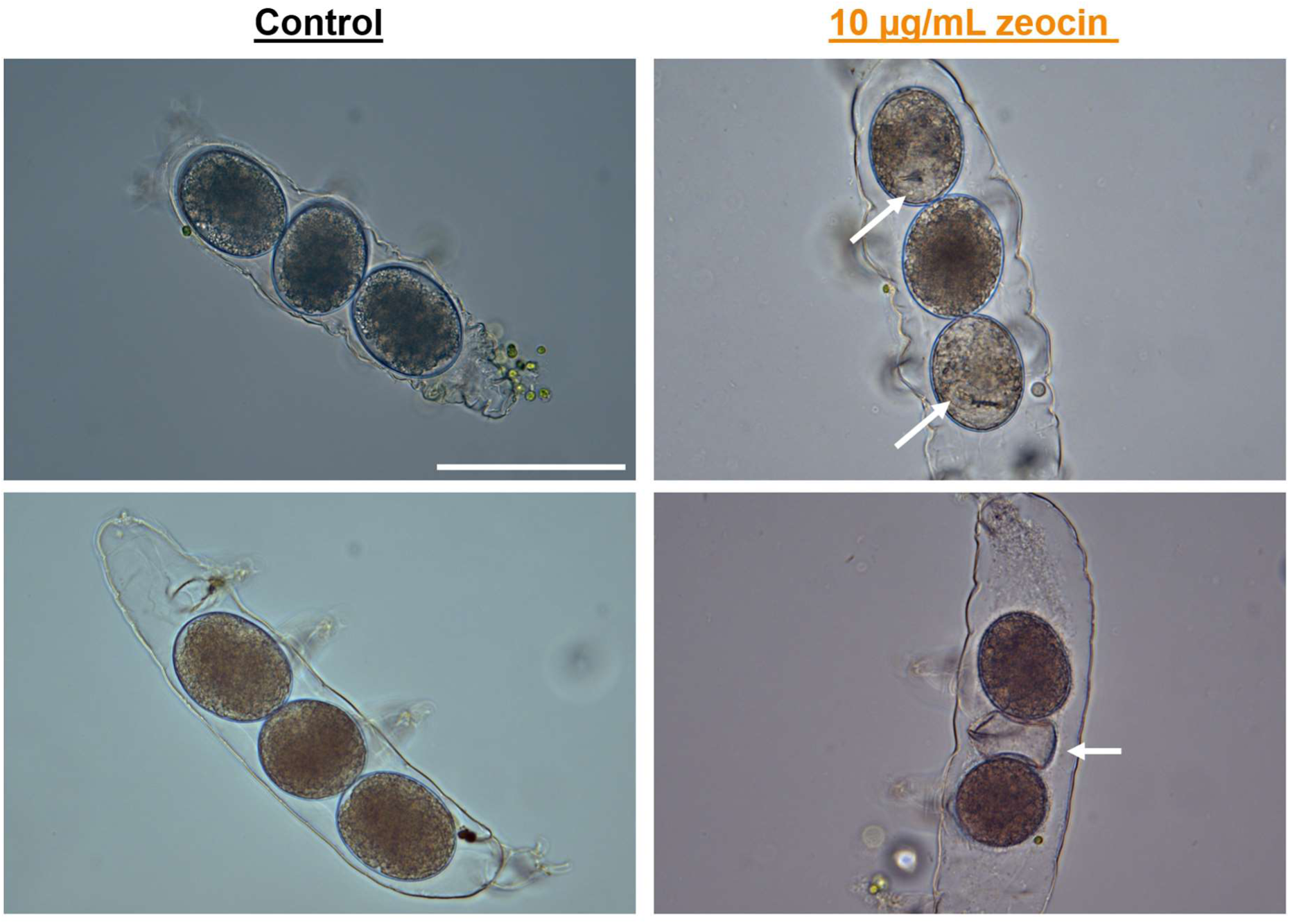
Exposure to 10 μg/mL zeocin induces embryonic development asynchrony. Representative images of exuviae containing developing embryos from control animals (left), showing synchronous development, and from animals treated with 10 μg/mL zeocin (right), displaying asynchronous development. White arrows indicate examples of asynchrony, such as the presence of a buccal apparatus in only 2 out of 3 embryos within the same exuvia (top right), or a hatched eggshell alongside two developing embryos (bottom right). Scale bar = 100 μm.

**Figure S12.**
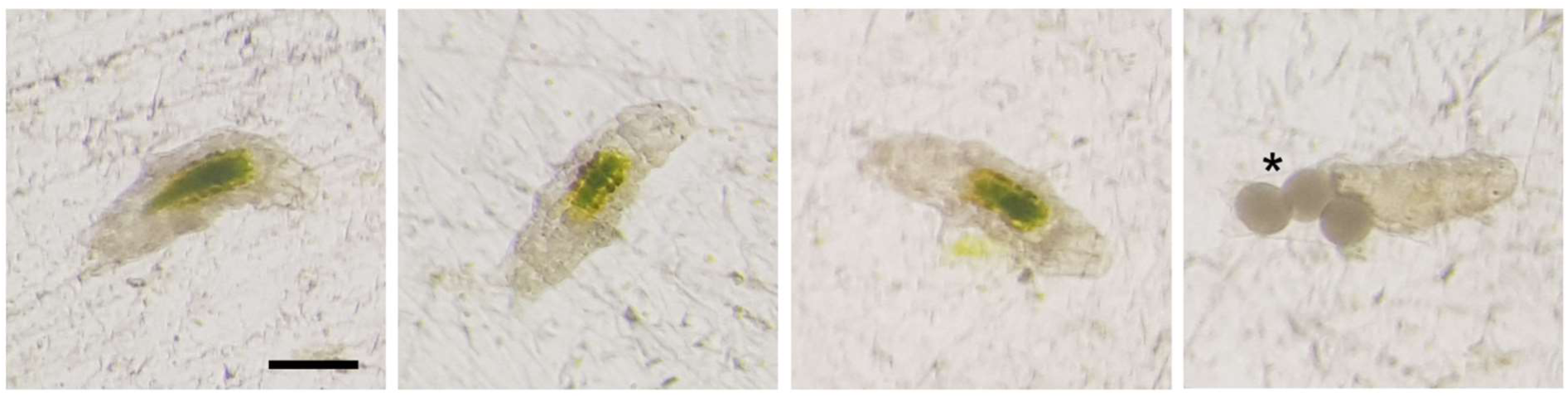
Embryos continuously exposed to 10 μg/mL zeocin develop into fertile adults. Representative images of adult *H. exemplaris* specimens continuously incubated in 10 μg/mL zeocin throughout embryogenesis and post-embryonic development. The animals exhibit normal feeding behavior, as indicated by algae-filled guts, and are capable of producing and laying eggs (asterisk), which successfully give rise to healthy, fertile second-generation offspring (not shown). *n* > 30. Scale bar = 100 μm.

**Figure S13.**
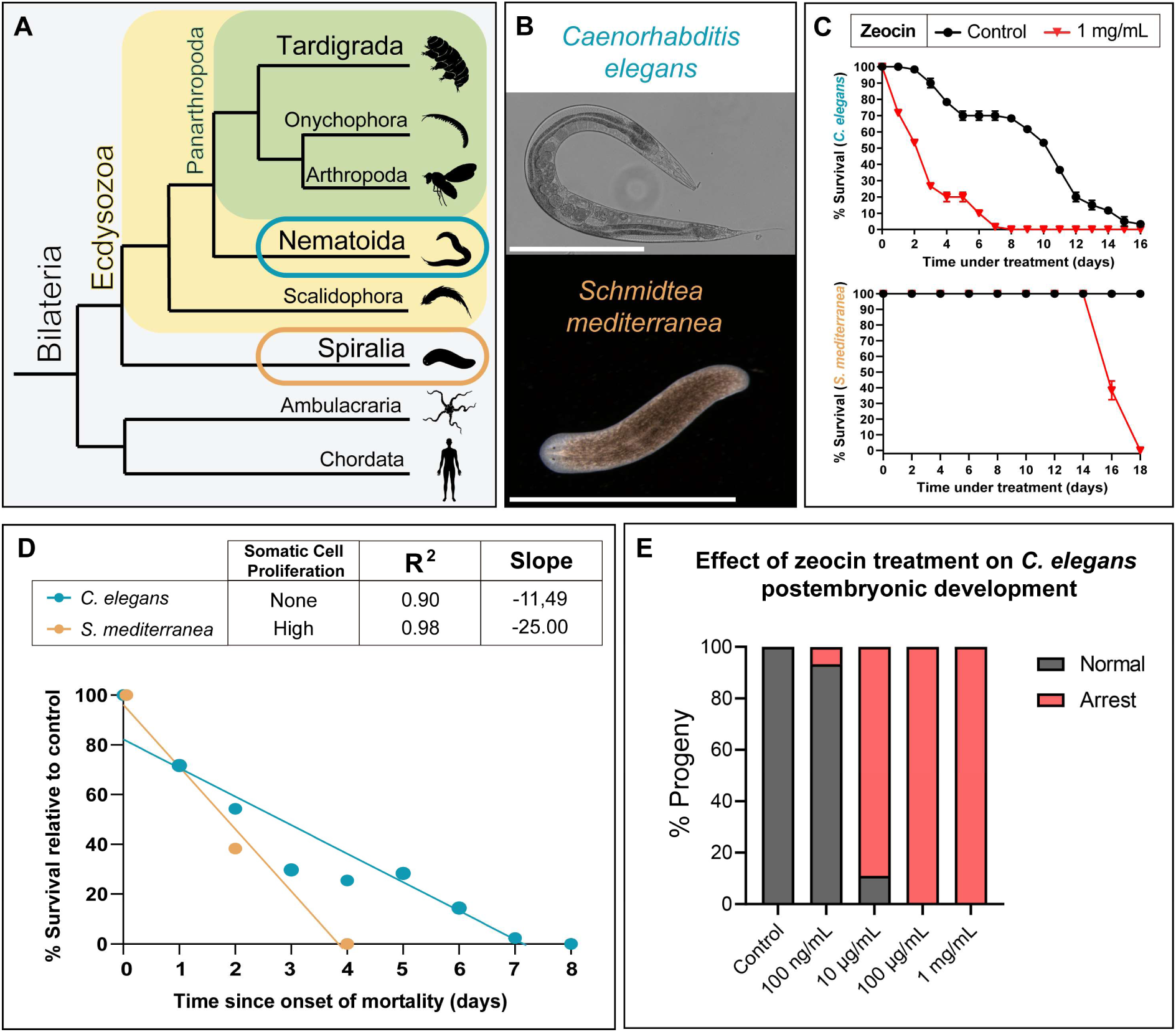
DNA damage–induced mortality correlates with proliferative activity across animal phyla. (A) Simplified bilaterian phylogeny highlighting the two species analyzed: *Caenorhabditis elegans* (Nematoida, within Ecdysozoa; teal blue) and *Schmidtea mediterranea* (Platyhelminthes, within Spiralia; orange-brown). Xenacoelomorpha are omitted for clarity. Images sourced from PhyloPic.org. (B) Representative images of *C. elegans* (Scale bar = 400 μm) and *S. mediterranea* (Scale bar = 6 mm) used in the comparative assays. (C) Top: Survival curves of adult *C. elegans* under control conditions and 1 mg/mL zeocin. Bottom: Survival curves of adult *S. mediterranea* under the same conditions. Data are shown as mean ± SEM from three biological replicates (*n* = 20 animals per replicate; 60 total in both species). (D) Survival regression curves for both species under 1 mg/mL zeocin, normalized to controls and aligned by time since the onset of mortality. (E) Histogram showing the effect of zeocin on *C. elegans* postembryonic development. The percentage of larvae in the progeny that arrested growth and died is plotted. *n* = 15-82 larvae per replicate depending on zeocin dose (at higher doses, worms produce less viable eggs). Values represent means from 3 biological replicates.

