## Supplementary Figures S1-S13 for "Compromised DNA replication in gut cells underlies tardigrade sensitivity to genotoxic stress"

*Running Title: Replication is a key vulnerability in tardigrades*

Key words: Tardigrade, Zeocin, Genotoxic stress, Gut cells, DNA replication

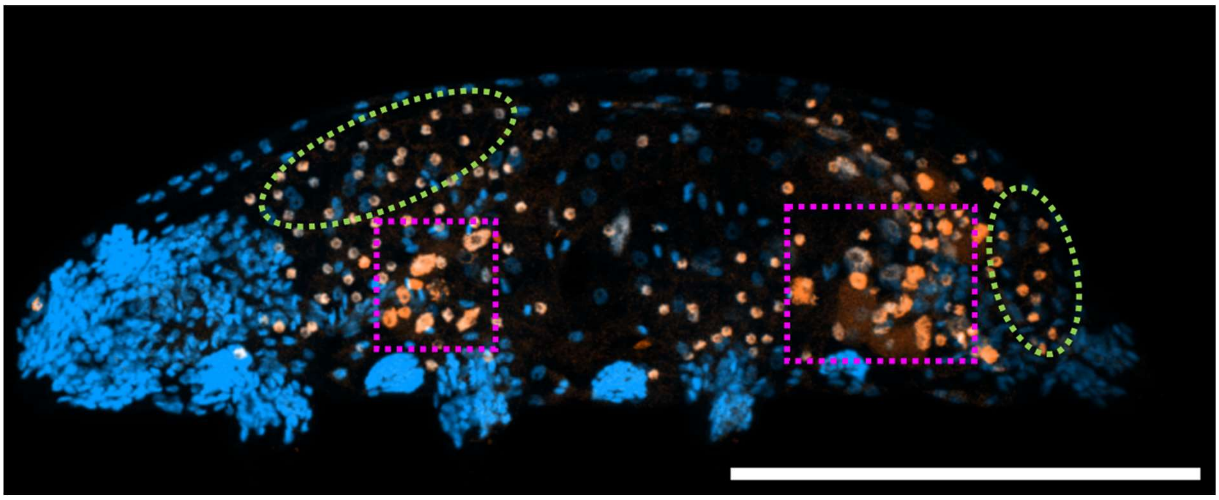

**Figure S1. Two-week EdU incubation reveals somatic proliferative cell types in tardigrades.** Representative maximum projection image of a tardigrade following a two-week EdU exposure. Nuclei are labeled with DAPI (blue), and proliferative ($EdU^+$) cells are shown in orange. Green circles highlight $EdU^+$ storage cells located in both anterior and posterior regions, while magenta squares indicate $EdU^+$ cells in the foregut and hindgut. Scale bar = 100 µm.

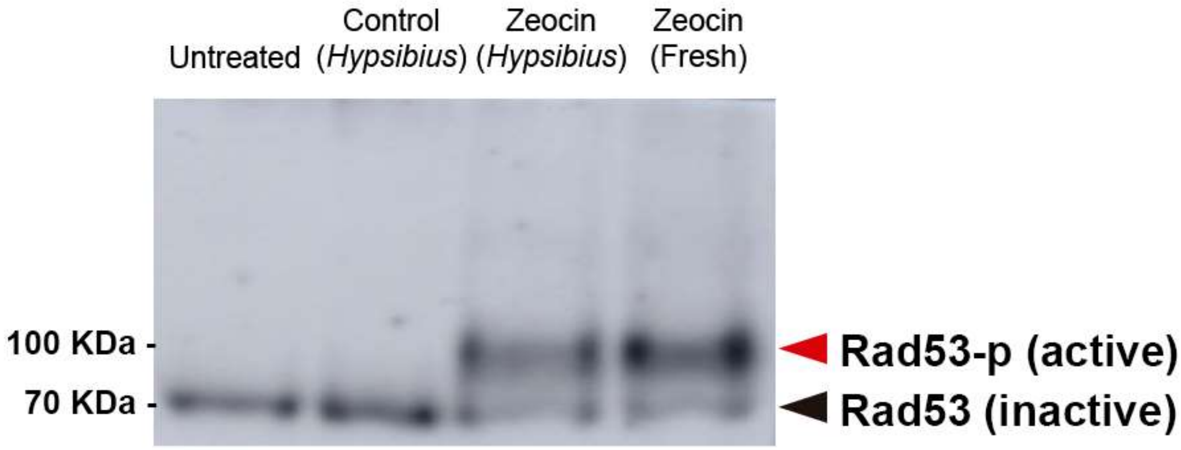


**Figure S2. Zeocin retains genotoxic activity in tardigrade culture medium for at least seven days.** Western blot of phosphorylated Rad53 (Rad53-p) in *S. cerevisiae* protein extracts. Black arrow represents Rad53 and red arrow represents Rad53-p, indicating an activation of the DNA Damage Response (DDR). Untreated = untreated yeast. Control (*Hypsibius*) = yeast treated with a 1/100 dilution of the water in which untreated *H. exemplaris* were incubated for one week. Zeocin (*Hypsibius*) = yeast treated with a 1/100 dilution of the 1 mg/mL condition in which *H. exemplaris* were incubated for a week [final concentration = 10 μg/mL]. Zeocin (Fresh) = yeast treated with fresh zeocin at 10 μg/mL. All yeast treatments were carried out for 3 hours prior to protein extraction.

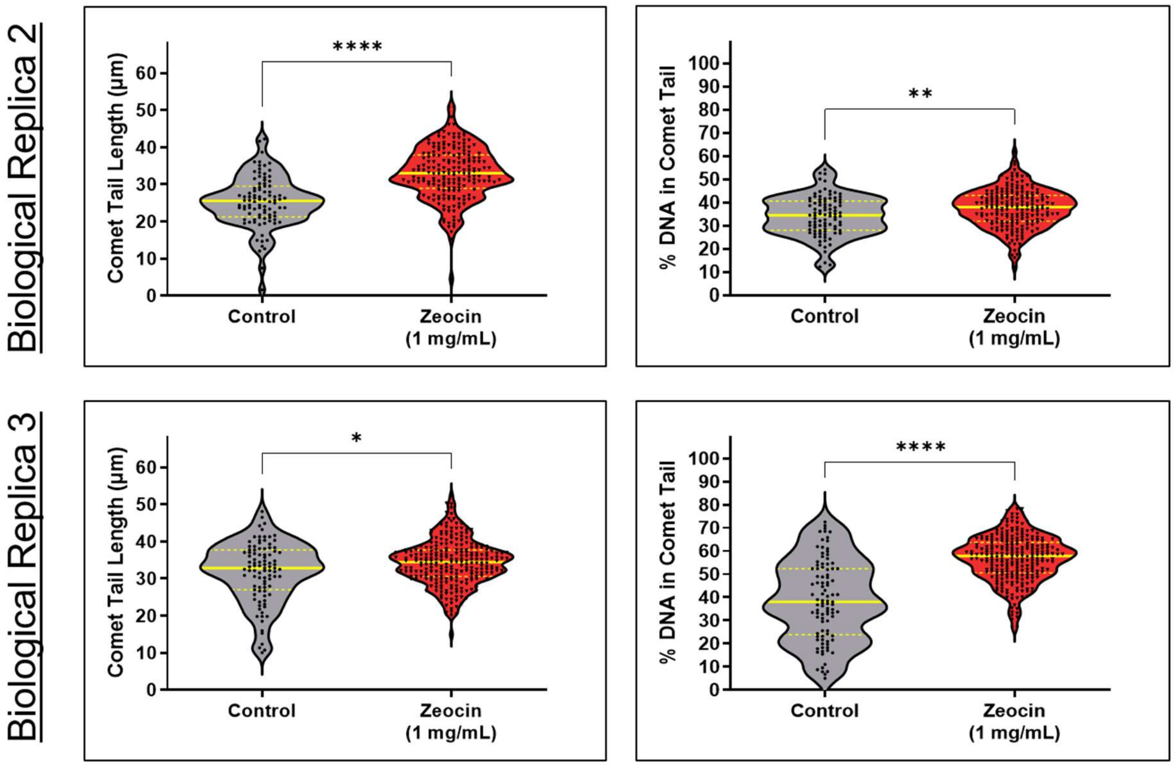


**Figure S3. Zeocin exposure induces DNA damage in tardigrade storage cells.** Violin plots showing comet tail length and percentage of DNA in the comet tail, comparing control and zeocin-treated (1 mg/mL for 4 days) storage cells from two biological replicates. In replicate 2, $n = 92$ (control) and $n = 192$ (zeocin-treated); in replicate 3, $n = 95$ (control) and $n = 255$ (zeocin-treated). Plot layout and quantification follow the format of Fig. 1F–G. **** $p \leq 0.0001$; ** $p \leq 0.01$; * $p \leq 0.05$ (Mann–Whitney U test).

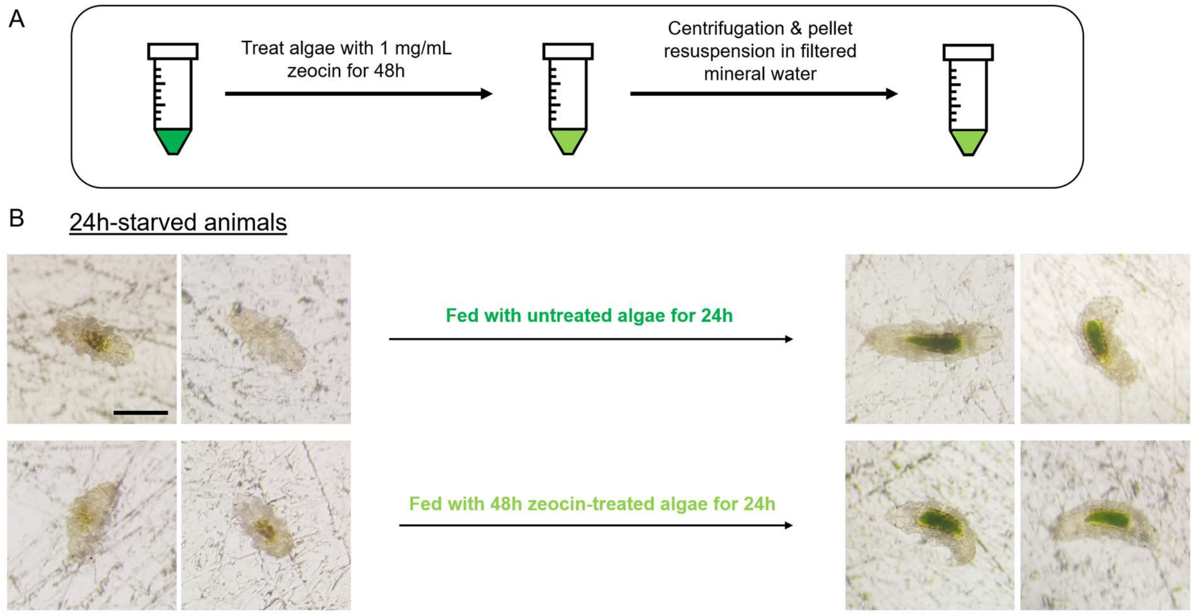


**Figure S4. Untreated *H. exemplaris* consume zeocin-treated algae.** (A) Schematic illustrating the treatment of *Chlorococcum* sp. algae with 1 mg/mL zeocin for 48 h, followed by centrifugation and resuspension in filtered mineral water for feeding to untreated tardigrades. The light green color denotes zeocin-treated algae, which show visible color decay after treatment. (B) Representative images of *H. exemplaris* specimens starved for 24 hours (left) and fed with either untreated or zeocin-treated algae (right). In both cases, the animals' guts are visibly filled with algae, indicating that untreated tardigrades are capable of ingesting zeocin-treated algae. $n$ > 30 in all cases. Scale bar = 100 µm.

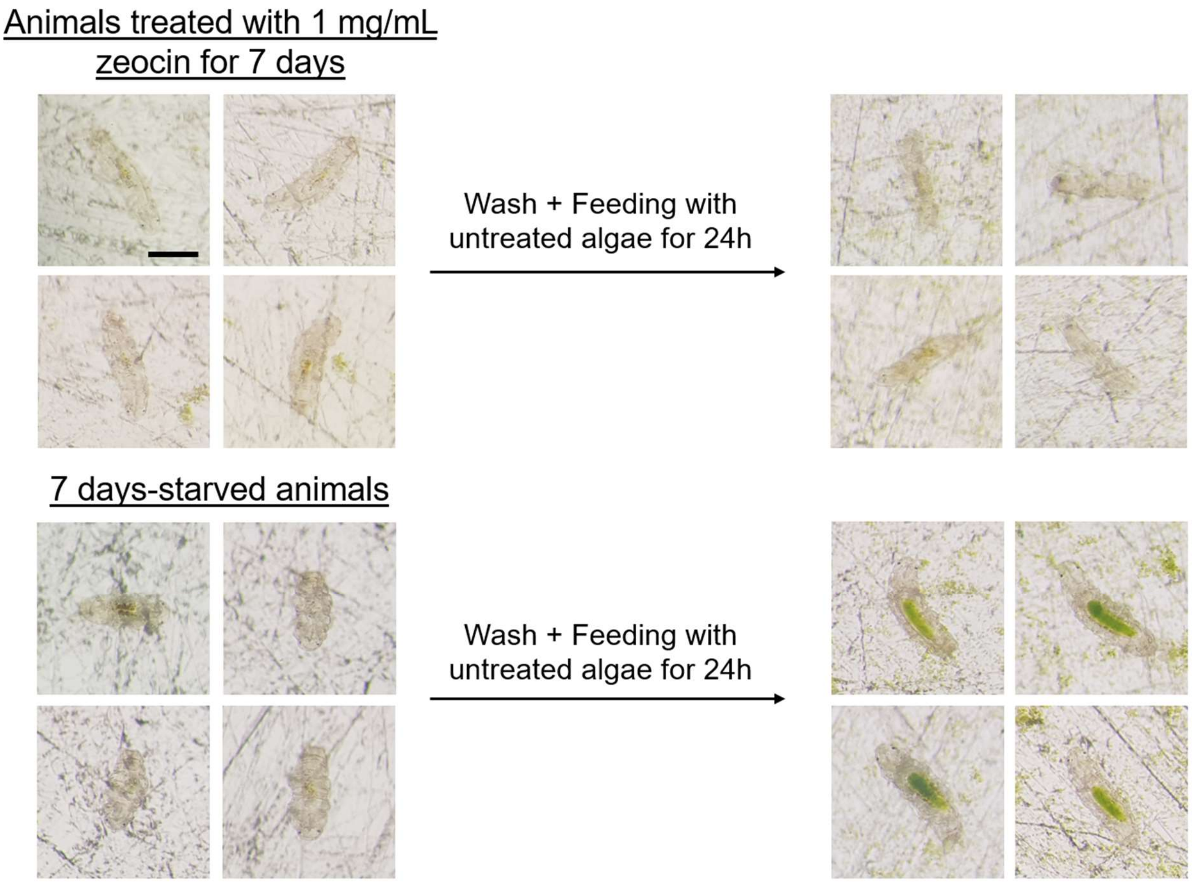


**Figure S5. Extended zeocin exposure impairs feeding in *H. exemplaris.*** Representative images of *H. exemplaris* specimens treated with 1 mg/mL zeocin for 7 days (top left), followed by 24 hours of feeding with untreated *Chlorococcum* sp. algae (top right). These animals show no visible algae in their guts, indicating a loss of feeding ability. In contrast, animals starved for 7 days without zeocin exposure (bottom left), then fed algae for 24 hours (bottom right), display guts filled with green algae, demonstrating preserved feeding capacity. *n* > 30 in all cases. Scale bar = 100 µm.

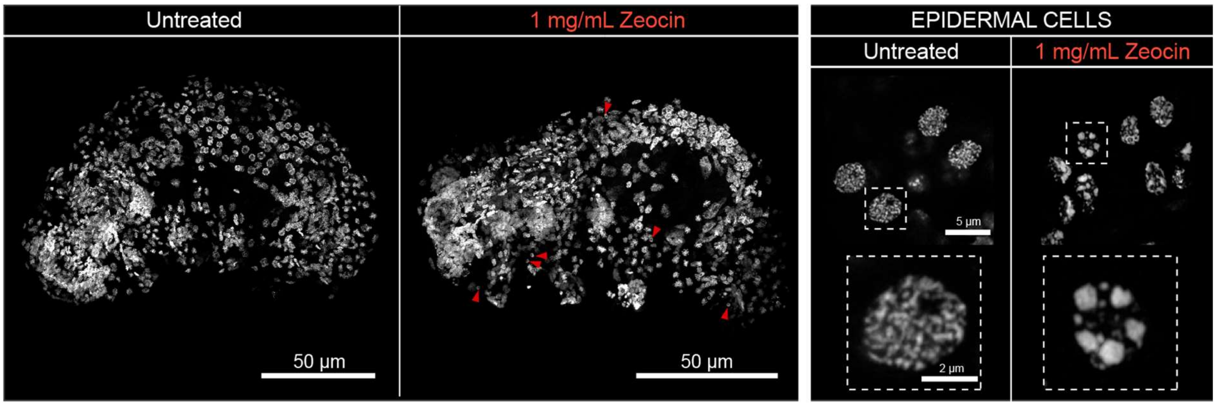


**Figure S6. Effects of zeocin on the DNA of *H. exemplaris* epidermal cells.** Left: Maximum projection images of whole animals stained with DAPI (nuclei, white) following a 1-week exposure to either untreated or 1 mg/mL zeocin conditions. Red arrowheads indicate affected nuclei in epidermal cells. Right: High-magnification images of epidermal cell nuclei from adult *H. exemplaris* under the same conditions, highlighting DNA fragmentation in zeocin-treated samples. Scale bars as indicated.

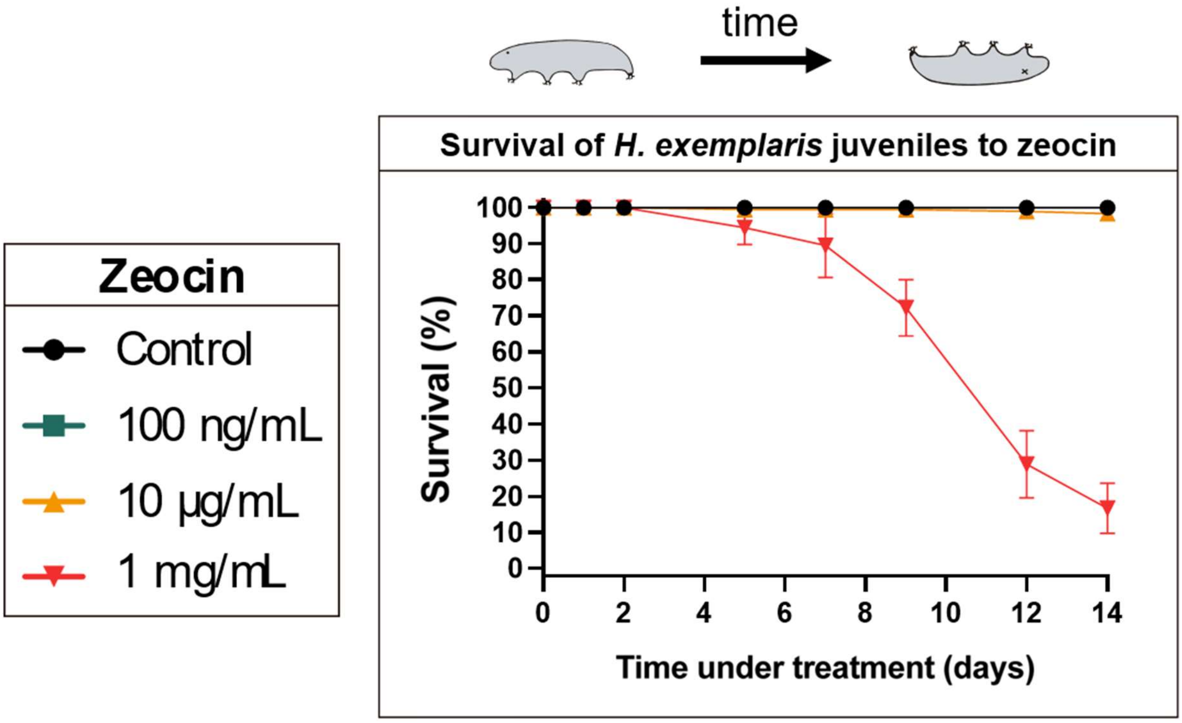


**Figure S7. Juveniles of *H. exemplaris* exhibit similar sensitivity to zeocin as adults.** Survival curves of *H. exemplaris* juveniles over time under continuous zeocin treatment at varying doses. Data are presented as the mean ± standard error of the mean (SEM) from three biological replicates, each consisting of 60 animals.

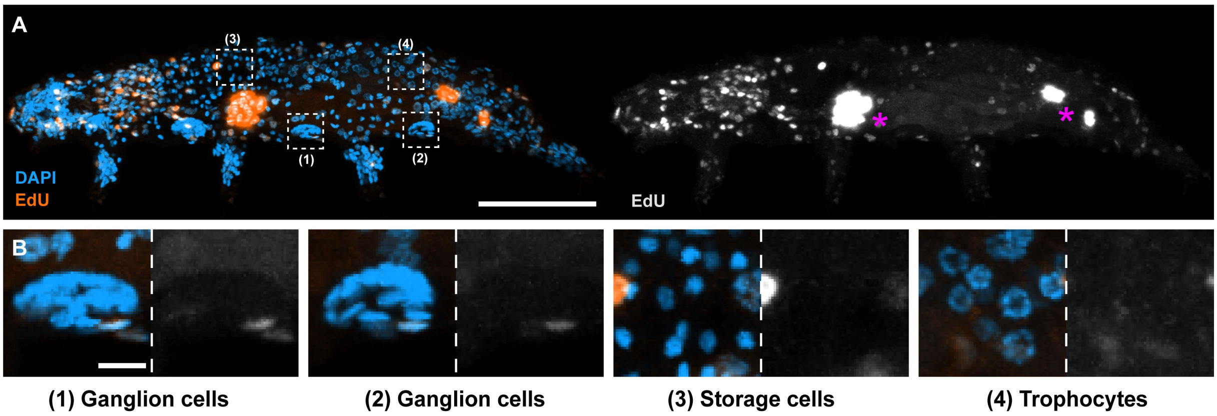


**Figure S8. Zeocin induces reparative DNA synthesis in select, but not all, cell types.** (A) Representative images of *H. exemplaris* after 4-day EdU + zeocin treatment, showing nuclei (DAPI, blue) and EdU$^+$ cells (orange). Right panel displays EdU signal alone (gray). Magenta asterisks mark EdU$^+$ gut cells in the foregut and hindgut. Dashed boxes highlight regions shown in (B). (B) Enlarged views of specific cell types from the boxed areas in (A). Note the absence of EdU incorporation in these cell types. Images are maximum intensity projections of full-depth z-stacks. Scale bars: 50 μm (A); 5 μm (B).

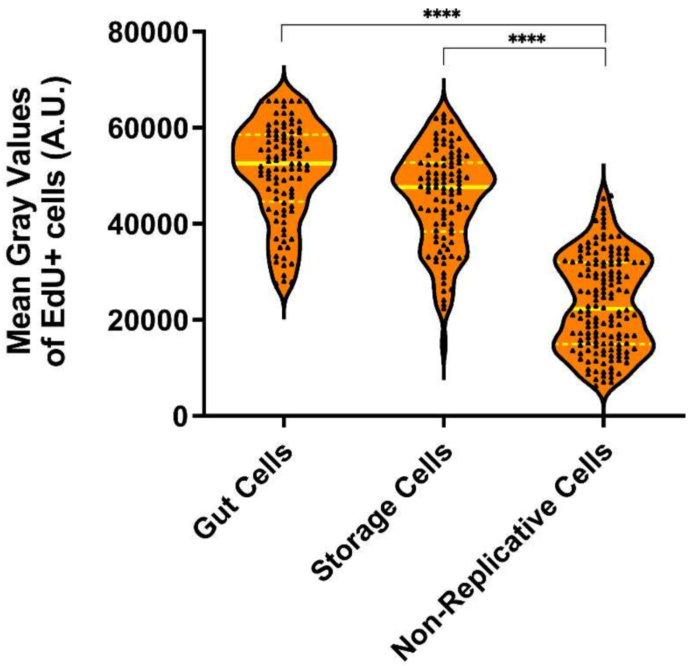


**Figure S9. EdU⁺ non-replicative cells display lower fluorescence levels.** Violin plots showing EdU fluorescence intensity in gut and storage cells (replicative), and in other somatic cell types (non-replicative), following zeocin exposure (as in Fig. 3A, C). Approximately 10 cells were quantified per animal from at least three animals across three independent experiments. $n$ = 100 (gut cells), $n$ = 101 (storage cells), and $n$ = 150 (non-replicative cells). Plot layout and quantification follow the format used in Fig. 1F–G. Each cell is represented by a triangle. A.U. = arbitrary units. **** $p \leq 0.0001$ (Kruskal–Wallis test).

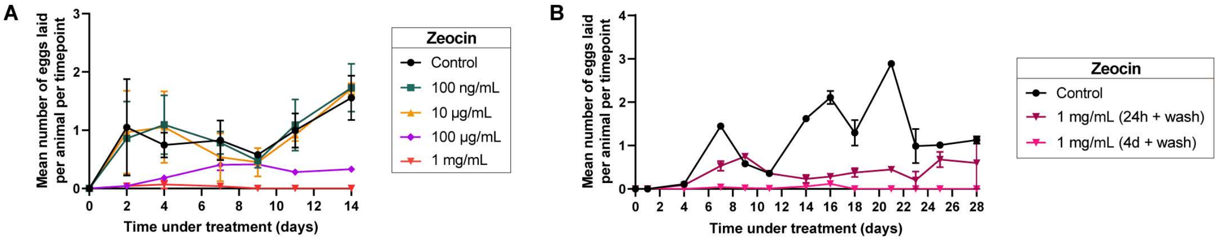


**Figure S10. Fertility of zeocin-treated *H. exemplaris* measured per timepoint.** (A) Average number of eggs laid per alive animal per timepoint over 14 days of continuous zeocin treatment at varying doses. Data are shown as mean ± standard error of the mean (SEM) from three biological replicates, each consisting of 60–120 animals. (B) Average number of eggs laid per alive animal per timepoint over 28 days following two pulsed zeocin treatment regimens (24 h pulse + wash; 4-day pulse + wash) at 1 mg/mL. Data are shown as mean ± SEM from two biological replicates, each consisting of 50 animals.

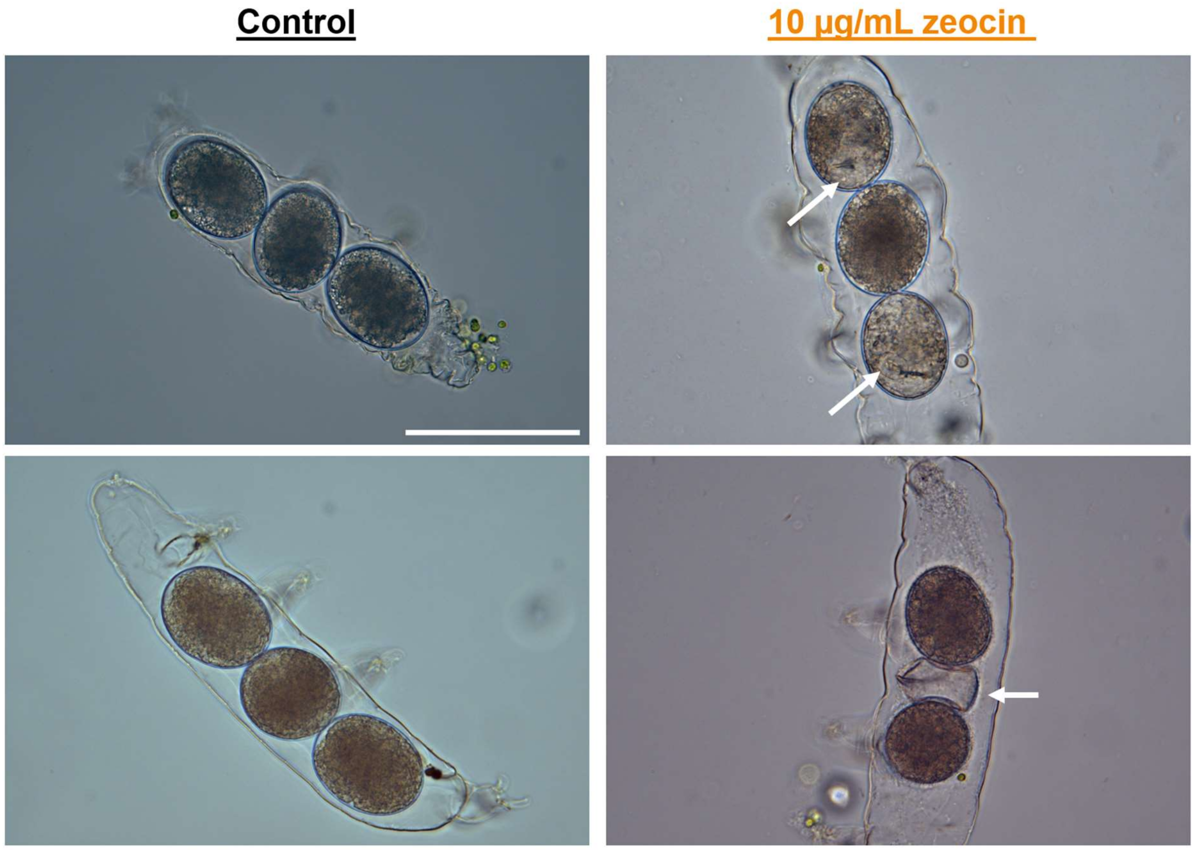


**Figure S11. Exposure to 10 µg/mL zeocin induces embryonic development asynchrony.** Representative images of exuviae containing developing embryos from control animals (left), showing synchronous development, and from animals treated with 10 µg/mL zeocin (right), displaying asynchronous development. White arrows indicate examples of asynchrony, such as the presence of a buccal apparatus in only 2 out of 3 embryos within the same exuvia (top right), or a hatched eggshell alongside two developing embryos (bottom right). Scale bar = 100 µm.

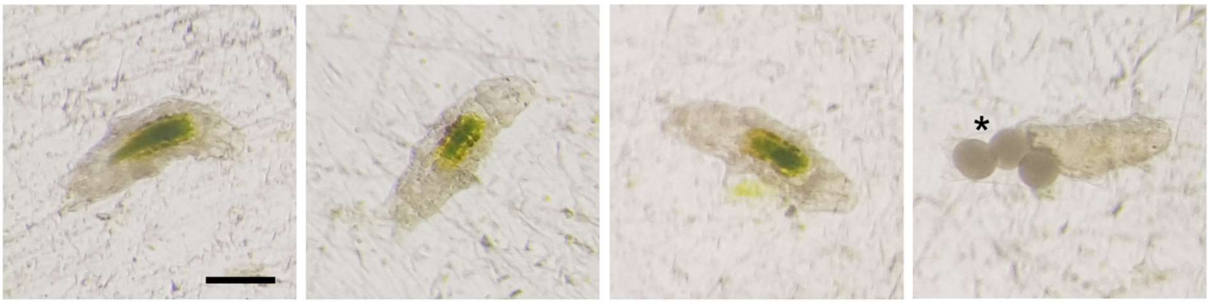


**Figure S12. Embryos continuously exposed to 10 µg/mL zeocin develop into fertile adults.** Representative images of adult *H. exemplaris* specimens continuously incubated in 10 µg/mL zeocin throughout embryogenesis and post-embryonic development. The animals exhibit normal feeding behavior, as indicated by algae-filled guts, and are capable of producing and laying eggs (asterisk), which successfully give rise to healthy, fertile second-generation offspring (not shown). $n$ > 30. Scale bar = 100 µm.

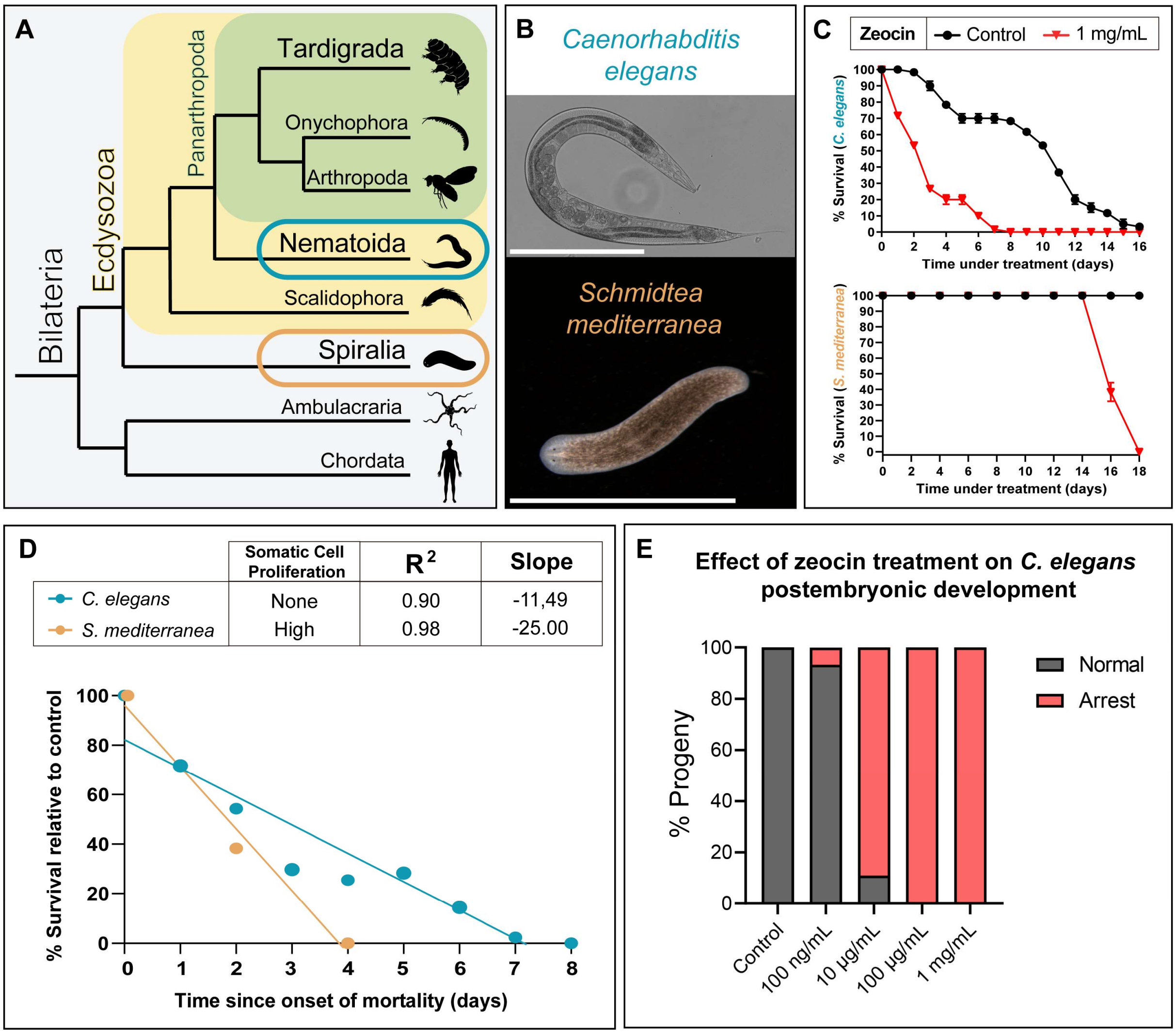


**Figure S13. DNA damage–induced mortality correlates with proliferative activity across animal phyla.** (A) Simplified bilaterian phylogeny highlighting the two species analyzed: *Caenorhabditis elegans* (Nematoida, within Ecdysozoa; teal blue) and *Schmidtea mediterranea* (Platyhelminthes, within Spiralia; orange-brown). Xenacoelomorpha are omitted for clarity. Images sourced from PhyloPic.org. (B) Representative images of *C. elegans* (Scale bar = 400 µm) and *S. mediterranea* (Scale bar = 6 mm) used in the comparative assays. (C) Top: Survival curves of adult *C. elegans* under control conditions and 1 mg/mL zeocin. Bottom: Survival curves of adult *S. mediterranea* under the same conditions. Data are shown as mean ± SEM from three biological replicates (*n* = 20 animals per replicate; 60 total in both species). (D) Survival regression curves for both species under 1 mg/mL zeocin, normalized to controls and aligned by time since the onset of mortality. (E) Histogram showing the effect of zeocin on *C. elegans* postembryonic development. The percentage of larvae in the progeny that arrested growth and died is plotted. *n* = 15-82 larvae per replicate depending on zeocin dose (at higher doses, worms produce less viable eggs). Values represent means from 3 biological replicates.
