## Supplemental File 1 for "Compromised DNA replication in gut cells underlies tardigrade sensitivity to genotoxic stress"

### Comet Assay for Tardigrade Storage Cells

All solutions and equipment were prepared fresh or cooled as noted. Unless otherwise specified, all incubations were performed at room temperature (RT) in the dark.

---

##### Buffers and Agarose Preparations

- **Lysis solution (pH 10)**
  - 4 M NaCl (46.75 g in 200 mL $H_2O$)
  - 100 mM EDTA (5.84 g in 200 mL $H_2O$)
  - 10 mM Tris-HCl (0.242 g in 200 mL $H_2O$)
  - Adjust pH to 10 with NaOH pellets; filter through a 0.2 µm mesh. Store at RT (up to 6 months).
- **Alkaline electrophoresis buffer (pH 13)**
  - 300 mM NaOH (10.20 g in ~850 mL $H_2O$)
  - 1 mM EDTA, pH 8 (1.7 mL of 0.5 M EDTA stock in ~850 mL $H_2O$)
  - Cool at 4 °C before use; prepare fresh or the day before.
- **Tris neutralization buffer (0.4 M Tris-HCl, pH 7.5)**
  - 24.23 g Tris base in 500 mL $H_2O$; store at RT.
- **Low-melting-point agarose (LMA, 1% in 1x PBS)**
  - 0.1 g LMA in 10 mL 1x PBS; melt at 90–100 °C, then cool to 37 °C before use; store 1 mL aliquots at RT.
- **SYBR Gold staining solution**
  - 1 µL SYBR Gold (Invitrogen, S11494; 10 000x) in 10 mL TE buffer (10 mM Tris-HCl pH 7.5, 1 mM EDTA); store at 4 °C, protected from light.
- **Agarose-coated slides**
  1. Clean slides with ethanol and air-dry.
  2. Coat with 1% agarose in $H_2O$ (0.5 g in 50 mL) by soaking; drain excess and dry horizontally ≥2 h, then store over silica gel overnight.

---

##### Storage Cell Isolation and Embedding

1. **Cell collection**
   - Filter animals to remove algae, then transfer to 5% glucose-monohydrate solution in $H_2O$.
   - Under a stereomicroscope, gently dissect the animals using tweezers and allow storage cells to sediment.
   - Collect ~10 µL of concentrated storage cells with a fine glass pipette (we recommend to use a mouth pipette for precision).
2. **Embedding**
   - Mix 10 µL cells with 70 µL 37 °C LMA; pipette gently to avoid bubbles.
   - Spread ~80 µL of the cell–agarose mix onto a pre-coated slide in a circular patch.
   - Chill at 4 °C in the dark for ≥10 min to polymerize.

---

**Alkaline Comet Assay**

1. **Lysis**
    - Immerse slides in lysis solution supplemented with 1% Triton X-100 for 1 h at RT.
    - Rinse briefly in $H_2O$ for 10 min.
2. **DNA Unwinding & Electrophoresis**
    - Place slides in a tank adapted for comet assays (e.g., Comet Assay Electrophoresis System ES II) kept at 4 °C
    - Flood with chilled alkaline electrophoresis buffer; incubate 45 min at 4 °C to denature DNA.
    - Run electrophoresis at 14 V (≈300 mA) for 20 min in the same buffer at 4 °C.
3. **Neutralization & Fixation**
    - Rinse slides once in Tris neutralization buffer for 5 min at 4 °C, then rinse in $H_2O$ for 5 min at 4 °C.
    - Immerse slides in 100% ethanol for 5 min at 4 °C.
    - Air-dry at 37 °C for 10–15 min.
4. **Staining**
    - Apply ~70 µL SYBR Gold solution per slide for 30 min.
    - Rinse briefly with $H_2O$; dry completely at 37 °C (overnight if convenient).

---

**Image Acquisition and Analysis**

- Capture comets using a fluorescence microscope with a 20x objective.
- Quantify tail length and percent DNA in tail using Comet assay software (e.g., OpenComet or CometScore).
